# Genes ruler for genomes, Gnodes, measures assembly accuracy in animals and plants

**DOI:** 10.1101/2022.05.13.491861

**Authors:** Donald G. Gilbert

## Abstract

Gnodes is a Genome Depth Estimator for animal and plant genomes, also a genome size estimator. It calculates genome sizes based on DNA coverage of assemblies, using unique, conserved gene spans for its standard depth. Results of this tool match the independent measures from flow cytometry of genome size quite well in tests with plants and animals. Tests on a range of model and non-model animal and plant genome assemblies give reliable and accurate results, in contrast to less reliable K-mer histogram methods. The problem of half-sized assemblies of duplication-rich *Daphnia* is addressed. A 20-year old *Arabidopsis* genome discrepancy is resolved in favor of 157Mb as measured with flow-cytometry. Not all genome DNA samples contain a genome, examples and reasons for this are discussed. The T2T completed human genome assembly of 2022 is complete by Gnodes measures, with about 5% uncertainty. With full genome DNA, Gnodes measures within 10%, usually within 5%, of flow cytometry, indicating they are both measuring the same content. Public URL: http://eugenes.org/EvidentialGene/other/gnodes/

## Introduction

Genome reconstruction is a Goldilocks problem: answers are often too hot, or too cold, the just right solution takes effort to discriminate among these outcomes. When Goldilocks’ problem is that of distinguishing near-identical twins, as with duplicated genes and highly repetitive chromosome spans, solutions must draw on both intrinsic and extrinsic evidence. This problem can be large, as reported by Vertebrate Genomes Project members recently [Kim et al. 2021], who found “between 25 to 60% of the genes were either completely or partially missing in the previous assemblies” of several vertebrate genomes. This problem results from reliance on accurate chromosome assemblies to model genes, where inaccuracies are common. This author and others have earlier shown [Gilbert 2013] that gene set reconstruction from RNA sequences is generally more accurate than those modeled on chromosome assemblies. But the problem of distinguishing high-identity duplicated genes is common to both approaches. A solution to both may be possible: use of DNA samples to carefully measure copy numbers of genes and chromosome spans.

A primary objective of this work is to measure the contents of genomic DNA samples from animals and plants, and provide a genomically informative summary of those contents, including chromosome sub-span details. These measures help describe accuracy of a chromosome assembly, inform projects of genomic DNA sample needs, and/or for different assembly methods. Existing genome informatics tools address this objective in many ways, but it is hard to find a straight-forward, reliable and biologically meaningful measurement of major components of genomic DNA. Genomes and their information content are complex, and vary much within species as well as among related species, so it is a challenge to measure them accurately with confidence.

Introductory material on the genome informatics presented here are well covered by two papers that developed DNA read mapping analyses, Pucker 2019 and Pflug et al 2020, and by two that developed DNA read k-mer frequency analyses, Vuture et al 2017, and Sun et al 2017, in the context of genome size estimation. Biochemical methods of genome size estimation are well established, and are subject to measurement errors of various kinds, but are not known to be biased [Dolezel & Greilhuber 2010, Pellicer & Leitch 2019]. This paper uses those as benchmarks for DNA sequence estimation, flow cytometry is the best known. DNA sequence estimation methods are also subject to various measurement and parameter choice errors. There is a question of k-mer frequency analyses being biased against against full accounting of duplicated or repetitive portions of genomes.

Genome size estimation from DNA reads are based on simple, meaningful algebraic equations, *Cm* = L*N/ G of Lander & Waterman 1988, and *Ck* = N(L - k+1) / (G - L +1) of eq.2 Li and Waterman 2003, where G genome size in bases, L a fixed length of DNA reads, N number of DNA reads, Cm, Ck are coverage depth of reads over the genome, k mer length for tabulating all existing sequence content of reads.

*Ck* and *Cm* are both coverage depths but differ due to k, *Ck* ∼ k/L * *Cm*, and both can be measured from DNA reads, to estimate G genome size. These measures are valid for certain assumptions, including that DNA reads are unbiased, randomly and evenly distributed cuts from genomic DNA. *Cm* is measured by aligning all DNA reads to an assembled model of that genomic DNA (chromosomes or gene transcripts), tabulating coverage depth across the assembly, then calculating mean or median values with error bounds using gaussian or near-normal statistics. *Ck* is measured by finding the peak (mode) of the frequency distribution (or coverage depth) of all k-mer fragments of the DNA reads, without use of an assembly model. Statistics pertaining to a poisson distribution of k-mers and frequencies analyses apply to Ck measures.

In this paper, and the cited read-map methods, Cm is measured using unique conserved gene sequences (UCG). These are unique biological sequences that can be verified by phylogenetic conservation, and generally assembly well due to their uniqueness. Thus UCG sequences provide a standard coverage depth for genomic DNA samples, and with the assumptions of randomly and evenly distributed DNA reads, all other genomic portions will have the same depth in a properly assembled diploid chromosome set.

Though coverage has extreme values, careful choice of unique measured genome spans reduces extremes and results in near-normally distributed, small variance for the statistic of Cm. Biological exceptions are ploidy variants: extra and missing copies of chromosomes (i.e X/Y), multiple non-nuclear genomes such as chloroplasts. In practice, chromosome assemblies are often deficient in duplicated regions, with a greater depth of coverage. Regions of depth below UCG are either over-assembled, or DNA samples are biased with reduced content of some regions.

Assumptions and components of Gnodes measurements include

A. A nuclear chromosome set is sampled for DNA, using methods suitable to reconstructing those chromosomes. The chromosome set is presumably diploid, variations in ploidy among them will affect results, but as Gnodes produces a table of chromosome (or contig) coverage depths, those variations can be identified.
B. Genome sequencing of this set produces a constant depth of coverage. Some random variance in depth is not a problem, but biases in depth are, which various sequencing methods have.
C. Unique conserved genes (UCG) identify single-copy regions of genomes, and thus measure the baseline constant depth of coverage. The BUSCO set of calculated conserved genes is used here [Simao et al 2015]. As this has a known level of gene duplications (i.e. 5% in original sets), as well unique genes can have duplicated subsequences (exons), Gnodes measures uniqueness qualities to exclude those genes with duplicated, skewed or uneven coverage.
D. DNA reads are aligned to both UCG and chromosome sequences, and measure correlated depth of coverage of these assemblies. Under- and over-assembly of chromosome regions are identified by the ratio of depths. Gnodes tabulates multi-mapped reads explicitly, unlike related depth measures, e.g. “samtools depth”.
E. Evidence annotations, from transposons/mobile elements, simple repeats, all genes, pseudogenes, are mapped to chromosomes and compared with read depth for identifying contents of over-, under- or just-right assembly. Included are DNA-supported gene parts missing from chromosomes, and gene models that lack DNA evidence.

A main output of Gnodes is a measure of over- and under-assembly (xCopy) relative to the standard depth of unique conserved genes. A DNA depth deficit analysis provides a synopsis of missing matter in assemblies and gene copy numbers. Detailed of coverage and contents of chromosome spans, including plots, allow detection of regions of mis-assembly, both over- and under-assembly. Measures of genes include copy numbers and coverage qualities, including deviations (skewed coverage, partial duplication, low or zero coverage). Gene copy numbers are calculated for DNA sample and for chromosome locations; a difference in these indicates genes with too few or too many chromosome assembly locations.

Genome assemblers produce rather different results for the same DNA samples. With a measure of their accuracy from Gnodes, the accurate portions from different assemblies can be merged, much like methods used for removing heterozygotic assembly spans [eg Purge_Haplotigs of Rhie et al 2021, eg. Falcon-Unzip of Chin et al 2016].

## Results

Gnodes measures coverage depth of a coding gene set, with required inputs of a coding gene sequence set, one isoform per gene, and precision genomic DNA reads, short enough to map well to coding sequences. Chromosome assemblies of that gDNA are also usual inputs, but are not required.

DNA coverage depth is measured by mapping sampled DNA fragments to chromosome and gene sequence assemblies, recording all duplicate, unique and null mappings, and cross-mapping of fragments on genes and chromosomes. Resulting summaries of these include genes coverage depth, where identified unique conserved genes (UCG) provide the baseline of depth in DNA sample (Table R1.1). The values of coverage depth, fragment size and number provide estimates of genome size, for model and non-model insects, plants, vertebrates and Daphnia species (Table R1.2). These are compared with flow cytometry sizes, genome assembly sizes, and estimates of genome sizes from statistical methods of K-mer fragmentation of DNA (Figures R1.1, R1.2). Further analyses of DNA depth deficits, and excesses, in chromosome contents and gene copy numbers are results (Figures R2.1, R2.2, R2.3). Chromosome graphs are produced of DNA depth and major components, to illustrate where in assemblies, and which components, have large discrepancies in the expected baseline DNA depth (Figures R3.1, R3.2). The problem of undersized Daphnia genome assemblies, and right-sizing them with DNA depth analyses, is addressed (Figures R4.1, R5.1, R5.2, R5.3, R5.4). The 20-year old Arabidopsis genome size discrepancy is analyzed with DNA depth analyses, and resolved in favor of flow cytometry measures (Figure R6.1, Tables R6.1, R6.2, R6.3). Genome size estimates are recapped: with complete genomic DNA samples, Gnodes estimates are within 10%, usually within 5% of flow cytometry, the other methods have much wider ranges (Table R7.1). Not all genomic DNA contains a complete genome, due to sampling methods or biological manipulation (Table R8.1, Figure R8.1), so that measurement accuracy depends much on DNA sample accuracy. Finally, the T2T complete human genome assembly is analyzed, and found complete within about 5% uncertainty (Table R9.1). Supplemental table S5 includes public IDs of chromosome and gene assemblies reported here. DNA samples are given with public SRA IDs.

### R1: Genome size estimation with Gnodes

The gene CDS by DNA measurement will estimate genome size, using formula G = L*N/C [Lander & Waterman 1988] for reads length L, reads number N, and DNA sample coverage depth C. C is measured by mapping DNA to unique conserved gene coding sequences. A genes DNA coverage table is Gnodes first primary result, with an example in Table R1.1.

The Gnodes algorithm for genome size measurements agree closely with those measured by the molecular method of flow cytometry (Figure R1.1). As these two methods use different evidence and techniques, the agreement suggests both are accurately measuring the same biological value. Current genome size measures based on k-mer counting have notably greater error in agreement with flow cytometry values. Chromosome assemblies also are typically below FC measured sizes, a well-known result [Peona et al 2018].

**Table R1.1.**
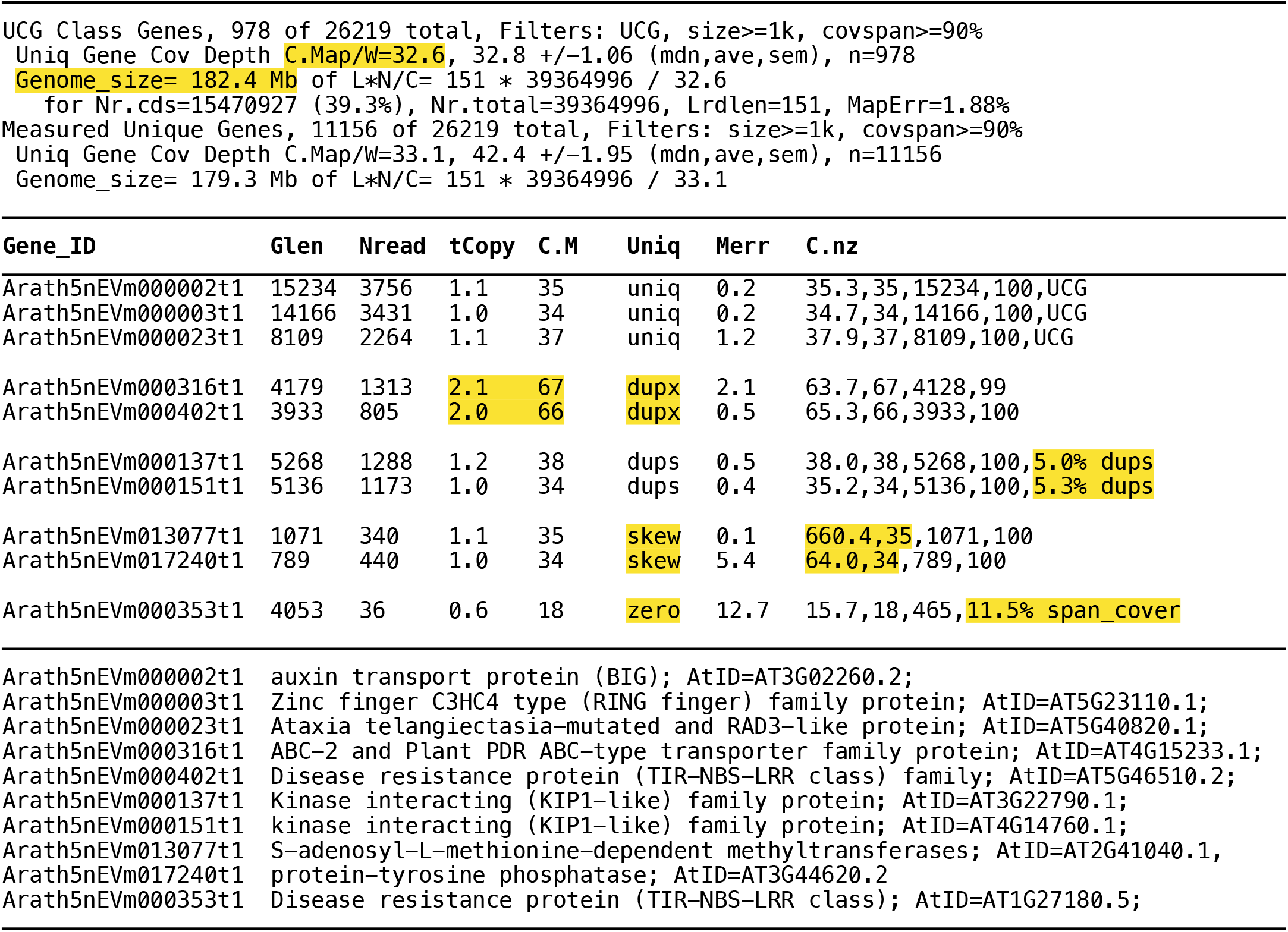
Gnodes Genes DNA coverage table example of *Arabidopsis thaliana* (arath17evg gene set, SRR10178325 DNA reads). The Uniq class column identifies 4 types: uniq, dupx (duplicated), dups (partial duplicate), skew (uneven coverage), and zero (coverage below reliability). Columns Glen, Nread are length of coding sequence, and number of reads mapped. tCopy is total copy estimate, C.M is coverage depth measure, Merr is map error percentage, C.nz is a tuple with average, median, length and percent covered span. The table top summary has unique gene cover depth measures, and genome size estimate from those and the read length and number in DNA sample (full table in Supplement S1).

**Figure R1.1.**
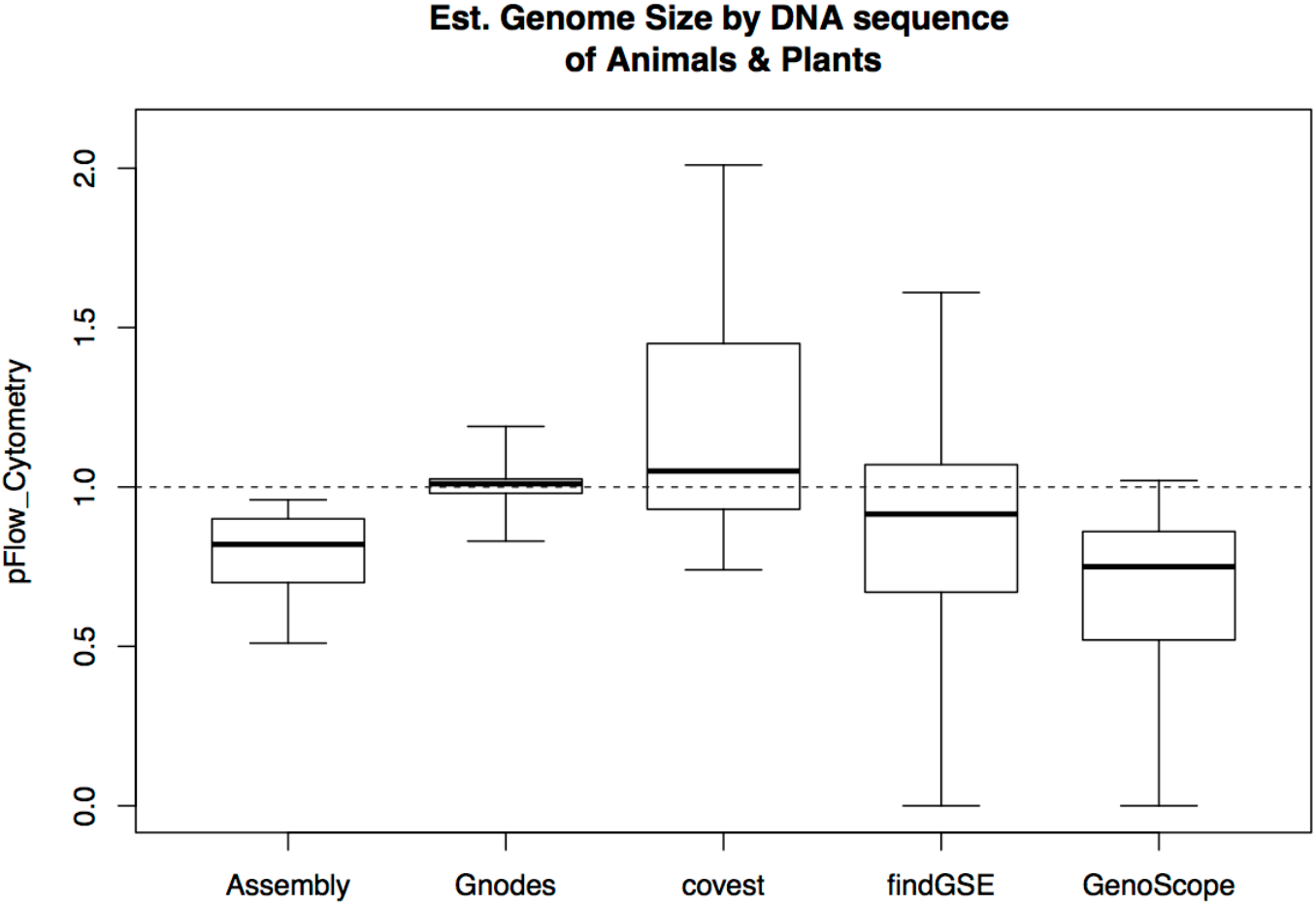
Boxplots (median, range) of estimators’ equivalence to flow cytometry (FC) measured genome sizes. Gnodes is very accurate, whereas k-mer histogram methods (GenomeScope, covest, findGSE) are rather inaccurate, with a wide range of estimates. Assembly sizes are typically below FC measured sizes. Detailed results of these size measures for 13 species are in Supplemental results.

GenomeScope [Vurture et al 2017], a widely used method, findGSE [Sun et al 2017], and covest [Hozza et al 2015] are K-mer based methods that are compared for genome size estimates. For the K-mer methods, two K-mer values were used, 19 (a common recommended default) and 29. Generally two gDNA samples per species, from different genome projects, were assessed. The DNA sample approx. cover depth ranged from 10 to 180, with read lengths from 100 to 250, for Illumina sampled DNA reads.

Gnodes will produce fairly reliable estimates at 10x coverage of precision short-read DNA, though 20x to 50x are recommended, while greater depths appear to add no reliability. Long reads (PacBio, Nanopore) can be used with Gnodes. They have two measurement problems for this use: high sequence error rates well above the level of gene and other duplications, and mapping long DNA to shorter coding sequences complicate depth measurement. Supplemental table S7 compares effects of different read-map software, which with default options have about 0.5% difference in number of reads mapped, with correlated effects on map error rate. Resulting size estimates differ due to map software by about 1%, but without clear correlation or suggestion of a superior method. Options available in mapping software will erase such discrepancies, by increasing or reducing number and precision of alignments. For long reads, minimap is used with Gnodes, or long reads can be cut to shorter pieces (eg. 300 bp longer than K-mer sizes of 20-30 bp) to better measure coding sequence depth (Table R8.1 includes long read measures).

GenomeScope and findGSE methods failed to produce estimates for some samples and K-mer values (recorded as zero estimates, Figure R1.1). Failures appear related to species genome contents (daphnia, fig, zebrafish and human) rather than other data parameters.

One serious flaw of all K-mer methods is that required choice of K read shred size strongly affects resulting estimates, and there is no biological basis for choice of K. These methods have found default or recommended K values that will produce size estimates to match some common species genomes (assembly or other measured size). However, species genome contents vary widely, affecting such estimates, such that K-mer choices for a given species have no firm basis. In contrast, unique conserved genes have a biological basis for providing a reliable DNA coverage depth: these are known single copy genes, and the resulting C measures for these have low variation, with a near normal distribution (median ∼ average ∼ modal values), indicating they measure a common value.

**Figure R1.2.**
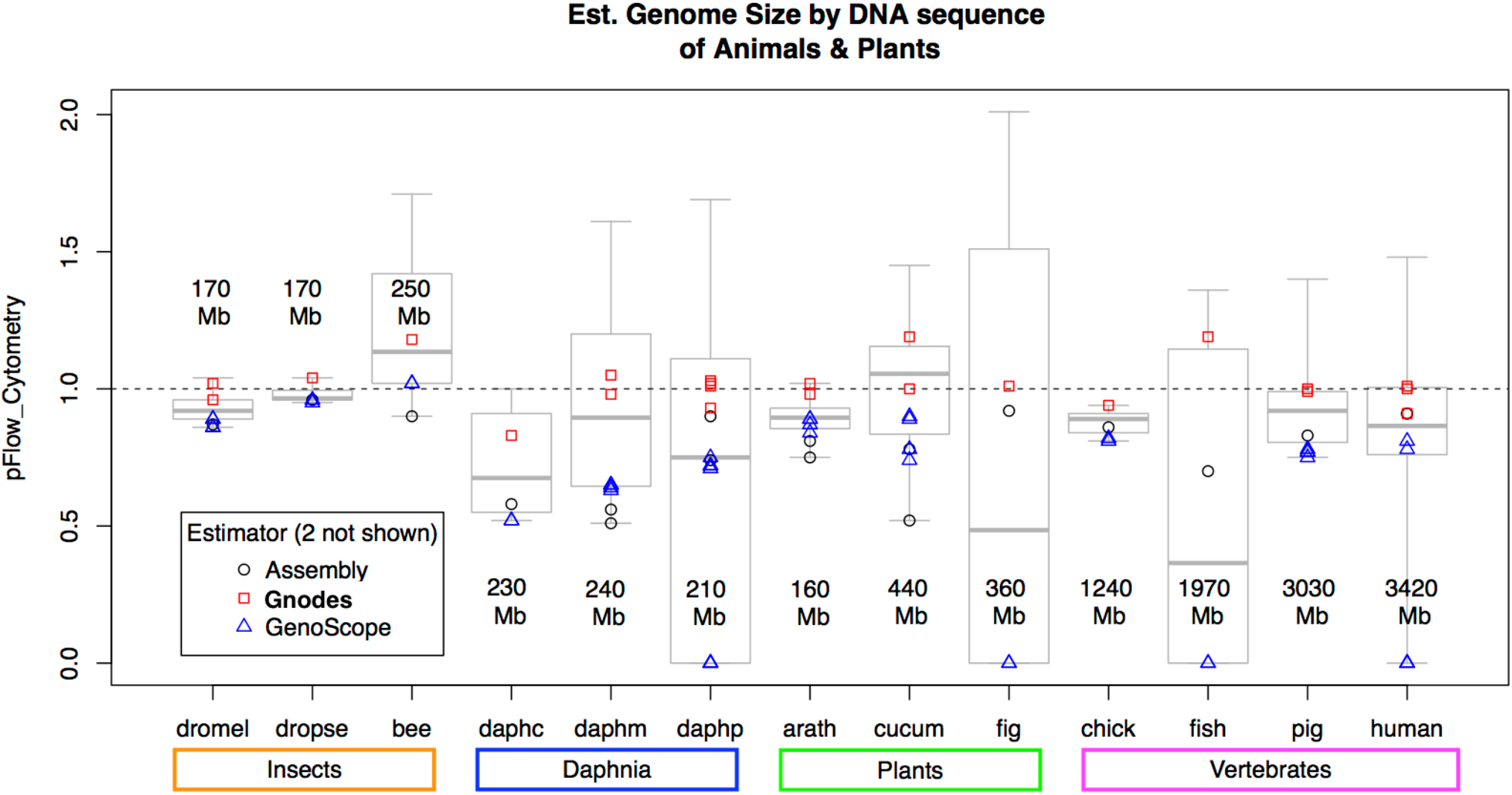
Estimations relative to FC value, for measured animal and plant genomes, with median, range and values from three estimators: Assembly, Gnodes, and GenomeScope. Flow cytometry sizes in megabases are given, ranging from 160 Mb (plant) to 3400 Mb (human). Detailed results of these size measures for 13 species are in supplemental results.

This agreement of Gnodes with flow cytometry extends over the range of small and large genomes of animals and plants, for well-studied model and less studied non-model organisms (Figure R1.2). There are variations in agreement, which likely depend on several factors including species-specific genomics and DNA sequencing methodology. Among two model species, Drosophila and Arabidopsis, certain DNA samples gave exceptional results (eg, heterozygous hybrids, samples from mutation or special populations). DNA samples that were sources of the chromosome assemblies proved most reliable, as these are procured with the intent of full assembly. Widest variation in measurements were found for zebrafish, fig tree, honeybee and daphnia water fleas, where species genome attributes likely are complicating such measures.

Gnodes assembly summary tables list observed and estimated sizes of annotated partitions of an assembly, as example Table R1.2 for *Drosophila melanogaster* assemblies.

**Table R1.2.** Gnodes assembly partition size summary for two assemblies of *Drosophila mel*., with flow cytometry size of 156-176 Mb. Obs is observed size (Mb), Est is estimated size from UCG cover depth, xCopy is a ratio of Est/Obs. Partition items: uniq, dup asm distinguished by duplicated read mapping, sum to all measured assembly; cdsasm contains reads also mapped to gene CDS; annotations CDSann, TEann, RPTann contain alignments of those sequence types (coding sequence, transposons and simple repeats), NOann lacks any of those.

| Item | DrosMel14r6 |  |  | DrosMel20Pi |  |  | Description |
| --- | --- | --- | --- | --- | --- | --- | --- |
|  | Obs | Est | xCopy | Obs | Est | xCopy |  |
| allasm | 138 | 165 | 1.20 | 163 | 165 | 1.01 | measured assembly |
| _total | 143 | 170 | . | 167 | 169 | . | total assembly |
| uniqasm | 115 | 128 | 1.11 | 104 | 113 | 1.08 | asm with unique gDNA |
| dupasm | 23 | 37 | 1.64 | 59 | 52 | 0.88 | asm with multimap gDNA |
| cdsasm | 29 | 41 | 1.40 | 33 | 35 | 1.05 | asm with CDS-mapped gDNA |
| CDSann | 25 | 27 | 1.07 | 27 | 26 | 0.97 | asm with CDS annotations |
| TEann | 24 | 27 | 1.10 | 32 | 28 | 0.87 | asm with Transposons |
| RPTann | 7.3 | 21 | 2.81 | 15 | 20 | 1.40 | asm with simple Repeats |
| NOann | 82 | 92 | 1.12 | 90 | 91 | 1.01 | asm without annotations |

**DrosMel14r6:** Total span=143.5 Mb, covspan=139.3, gaps=1.1 Mb for assembly drosMel6ref\_chr Genome Size Est= 166.8–175.6 Mb (Nread), 161.4–169.8 Mb (Maprd), for readset SRR11460802, for Size=LN/C, Cov=32.3,34, N\_reads=45376973, N\_maprd=43888939,96.7%, L\_readlen=125 **DrosMel20Pi:** Total span=167.8 Mb, covspan=165.2, gaps=0 Mb for assembly drosMel20pi\_chr Genome Size Est= 162.1–175.6 Mb (Nread), 155.5–168.5 Mb (Maprd), for readset SRR11460802, for Size=LN/C, Cov=32.3,35, N\_reads=45376973, N\_maprd=43534114,95.9%, L\_readlen=125 Uniq Conserved Gene Cover: median=32.3, ave=32.4, sem=1.32, n=606

### R2: Gnodes DNA Depth Deficit Analyses: Assembly Components and Gene Copy Numbers

Best use of Gnodes analyses is to compare two or more assemblies, possibly with several DNA samples. To that end, a visual graph of deviations from expected whole genome values, of genes and chromosomes, is of value. A deficit/excess synopsis plot of major components produced by Gnodes is a graphic summary of two tabular summaries: chromosome contents, and gene copy numbers. These synopsis plots add to tabular results of following arabidopsis and human assembly comparisons.

**Deficit** is the difference of observed assembly and gene copy numbers, from expected contents based on uniform DNA coverage depth, as measured over unique conserved genes. Percent deficit is shown (Y-axis) as ratio of this difference: 100 * (obs - exp) / exp. The desired value of 0 indicates chromosome assembly has all expected DNA coverage depth. A value above 0 indicates excess, below zero is a deficit. Y-axis for Gene-CN on right, with minimum -10% deficit, is narrower than for Chr Assembly on left.

**Chromosome assembly components (gray)**, ALL: all assembly, UNIQ: unique read map spans, DUP: duplicate map spans, CDS: Gene CDS-mapped read spans. UNIQ, DUP are full partitions of ALL (UNIQ+DUP=ALL), but need careful interpretation: a deficit in UNIQ here means that duplicates are lower than expected (ie under-assembled duplications). Excess in DUP means some are unique (ie over-assembled). A deficit in one balanced by excess in the other means roughly that it is *just-right assembled*, e.g. *Drosophila mel*.2020 Pi assembly.

**Gene copy number levels (green)** C1: one copy in genomic DNA reads mapped to gene-cds, C2-9: two to nine copies, C10-99: ten to ninety-nine copies. These values are the percentage of genes in measurement set with a deficit in copies found on chromosome assemblies, using CDS-mapped genomic DNA reads. This doesn’t measure total of missing copies, but those of the measured set with a deficit.

**Cmiss (red)** : Gene CDS-mapped genomic DNA reads that are missing from Chr-assembly, as percent of all CDS-mapped reads. Measurable Cmiss percentages (>=1%) indicate a poor assembly, i.e. missing all copies of gene coding sequence DNA.

Gnodes produces summary plots of deficit/excess in chromosome assembly components and gene copy numbers. The following Figures R2.1, R2.2, R2.3 summarize those content deficits, with some excesses, for Arabidopsis, Drosophila, Human, Pig and Chicken genome assemblies.

**Figure R2.1.**
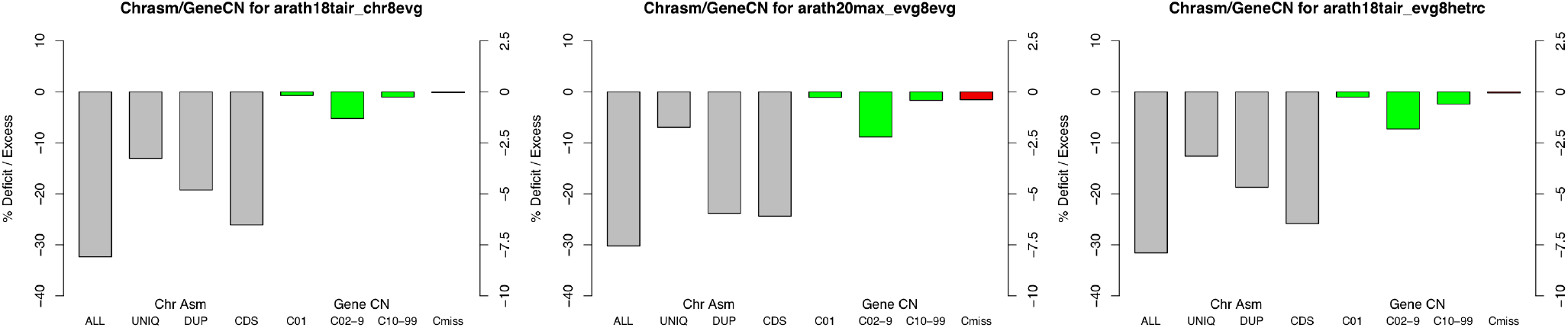
Gnodes Assembly & Gene Copy Deficit plot for *Arabidopsis thaliana*: Arath18TAIR and Arath20Max assemblies, and Arath18TAIR x F1 heterozygote DNA sample.

Arath18TAIR assembly has a -30% deficit in DNA spans, notably including genes with 2-9 copies. Arath20Max assembly is larger by 10 Mb or 6%, and has about 6% lower DNA deficits in chr assembly, notable it has 50% more spans with simple repeats. However this Arath20Max has a greater deficit in gene copy recovery, and 10x more missing unique gene DNA (0.40% Arath20Max vs 0.04% for Arath18TAIR), possibly an effect of sample population differences.

Heterozygous DNA (F1 of Col-0 x Cvi-0, tair_evg8hetrc panel) has no significant effect on genome-wide measures, versus homozygous DNA of Col-0 strain (tair_chr8evg panel). Measurable effects of F1-DNA are (a) higher map error rate, with more incomplete gene span aligns (28% F1 vs 5% parent Col-0) and (b) a small number of genes with copy number changes (10% with more or fewer copies in F1 mix).

Gene copy deficits in Arath18TAIR include Ribosomal proteins, Cytochromes, Transcription factors, Plant self-incompatibility, Disease resistance, Transmembrane genes, Transposon genes, among others. Uncharacterized genes account for roughly half of copy deficits.

**Figure R2.2.**
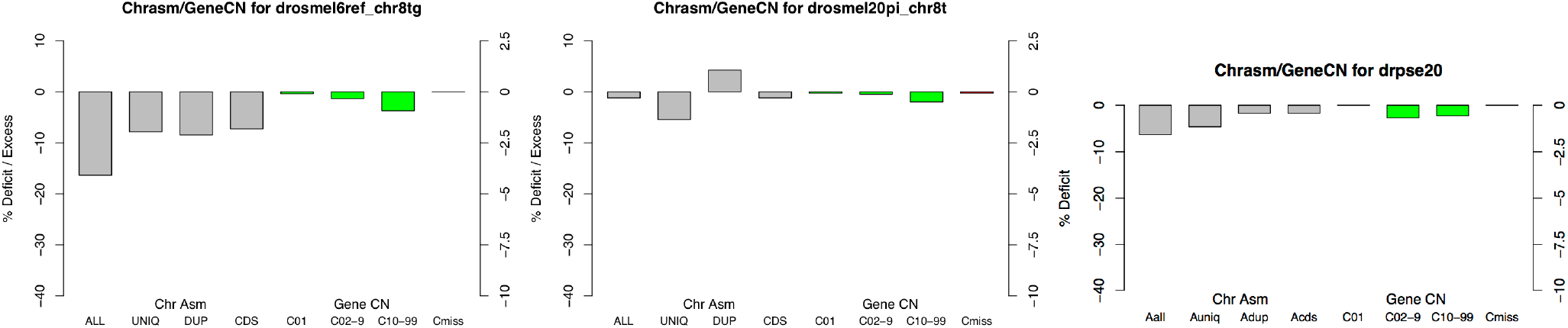
Gnodes Assembly, Gene Copy Deficit plot for *Drosophila melanogaster* drosmel6r, drosmel20pi and *Dr. pseudoobscura* 2020 assemblies.

*Dros. melanogaster* release 6 reference assembly of 2014 (drosmel6r) has a noticeable deficit of -17%, or 30 megabases. A more recent *Dros. mel*. 2020 (drosmel20pi, strain Pi2) and *Dros. pseudoobscura* 2020 (drospse20) assemblies are at zero deficit, within measurement error. The drosmel20pi apparent deficit Uniq + excess Dupl parts sum to zero deficit. The reference release 6 deficit is in part found among genes with 2-9 and 10-99 copies in genome, and also in transposon and repeat regions (not shown). Gene copy deficits in drosmel6r include Histones, Chorion, Mucin, Ubiquitin and uncharacterized genes among others. Histones have the majority of deficits. The drosmel20pi assembly reduces copy deficits for 20 histones, 4 mucins and 36 uncharacterized genes, though it has some deficits not in drosmel6r assembly, including male genes as it lacks ChrY. There are no transposon genes in the *Dros. melanogaster* gene set used for annotations.

**Figure R2.3.**
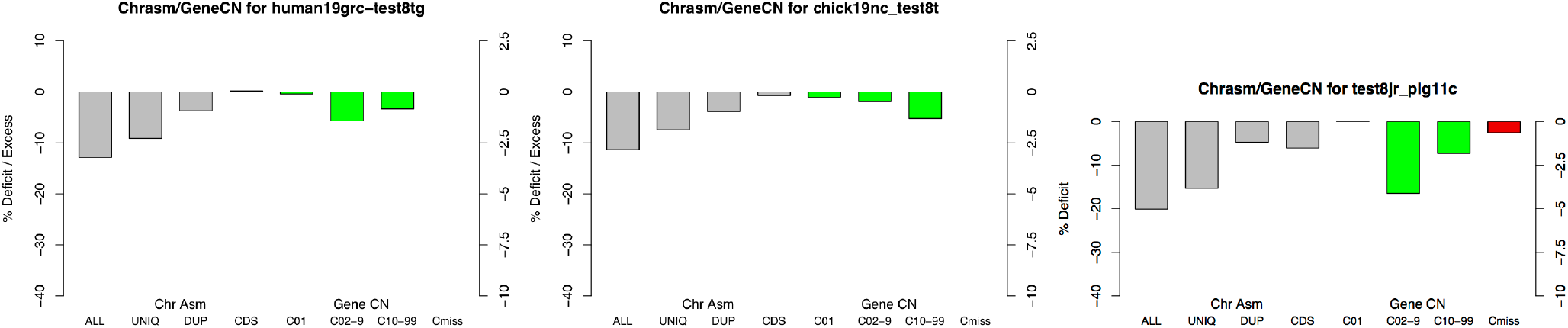
Gnodes Assembly, Gene Copy Deficit plot for Human, Chicken, Pig assemblies.

The reference human19grc assembly is close to accurately recovering flow cytometry measures of genomic DNA (3099 of 3423 Mb). Proportionally largest deficits are in simple repeat spans, of 50-80 Mb, and all duplicated spans including repeats, of 75-125 Mb. Gene copy number deficits for human18grc are largest for a small number of families with 10-99 copies, and a larger number of 2-9 copy genes missing one copy. Notable gene copy deficits are found for olfactory and taste receptor gene duplications, homeoboxes, antigens, ribosomal proteins, zinc finger containing and uncharacterized genes. Two human chromosome assemblies were measured, reference human19grc and more a recent human20ash (see below **R: Human genome assembly** for 2022 assembly). The later is 2% larger, has minimal differences measured by Gnodes from the reference assembly, in both whole genome partitions and in gene copy numbers, and is not shown here.

Chicken and pig reference chromosome assemblies are also analyzed with Gnodes. Pig genome size is similar to human, chicken is a third of that at 1200 Mb. Chicken appears to be close to accurately assembled. Pig however has significant deficits in whole chromosome parts, in gene copy numbers as well as missing unique gene DNA (missing 0.670% of pig gene DNA, vs 0.008% for human, 0.017% for chicken). Olfactory receptor gene duplications are an aspect of biological value that may be improved in pig’s chromosome assembly. Pig assembly has many fewer annotated olfactory genes than human, but also many more copy deficits in these (pig misses copies in 50 of 100 genes, versus human misses copies in 20 of 400). Chicken assembly shows copy deficits in about 20 of 270 olfactory receptor genes.

### R3: Gnodes DNA Depth Chromosome Plots: DNA-xCopy and Major Components

Whole genome measures and statistics are prone to mis-interpretations, as genome devils are in details. As many readers know, a “correct” whole genome measure can result from averaging many complex, incorrect but contrary details. An aid to proper interpretations is a visual plot of measured contents of chromosomes that can be zoomed in to problem spans. Gnodes produces chromosome plots of copy depth and major annotation components (coding sequence, transposon and simple repeat) on chromosomes (or scaffolds), in magnifiable PDF format. Figure R3.1 (Arabidopsis) and Figure R3.2 (Drosophila) show snippets of chromosome plot starts, with graph of XCOPY, DUP duplicated DNA, CDS coding gene, TE Transposon, and RPT simple repeats as percent of spans. XCOPY is DNA copy depth divided by standard of Unique Conserved Gene copy depth, with black bar at 1. Deficit (left) and Excess (right of 1-bar) depths are visible as red graph. Excess depths suggest span is **under-assembled**, seen most often for RPT and DUP regions, though mis-mapping of duplicate DNA may account for some discrepancy.

Arath18TAIR assembly has a -30% deficit in DNA spans, notably including genes with 2-9 copies. 2020Max assembly, though larger by 10 Mb, has greater deficits including large missing gene DNA. *Dros. melanogaster* release 6 (drosmel6r) has a noticeable deficit of -17%, or 30 Megabases. Recent *Dros. mel*. 2020 (drosmel20pi) and *Dros. pseudoobscura* 2020 (drospse20) assemblies are at zero deficit, within measurement error. In drosmel20pi, XCOPY variation on the chromosomes is higher than in drosmel6r, suggesting some errors in this new assembly. The drosmel6r assembly has many un-located scaffolds with a high XCOPY variation, not shown, that account for the noticeable deficit in assembly size.

**Figure R3.1.**
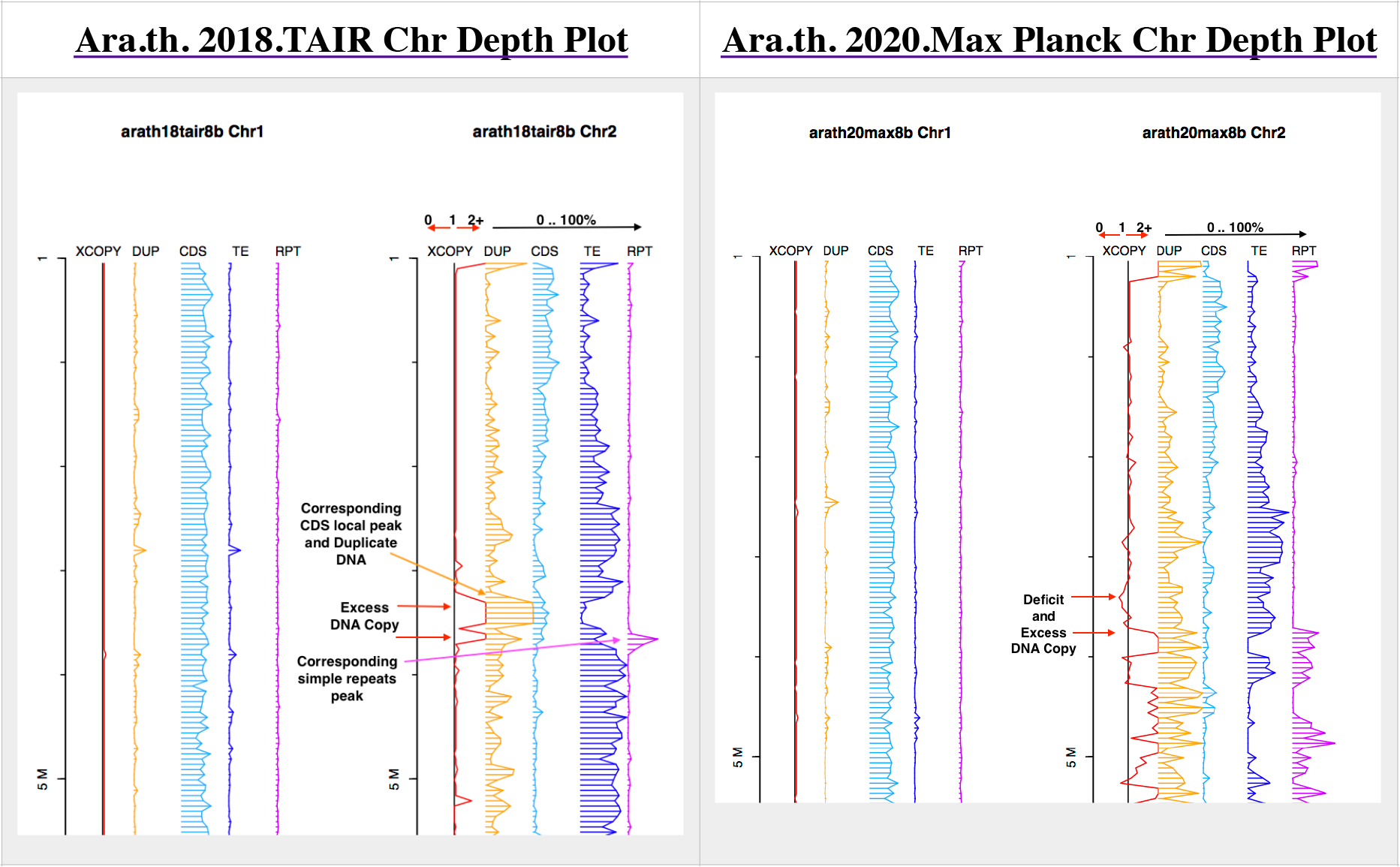
Gnodes Chromosome plot of *Arabidopsis thaliana*: arath18TAIR, and arath20Max Planck assemblies

**Figure R3.2.**
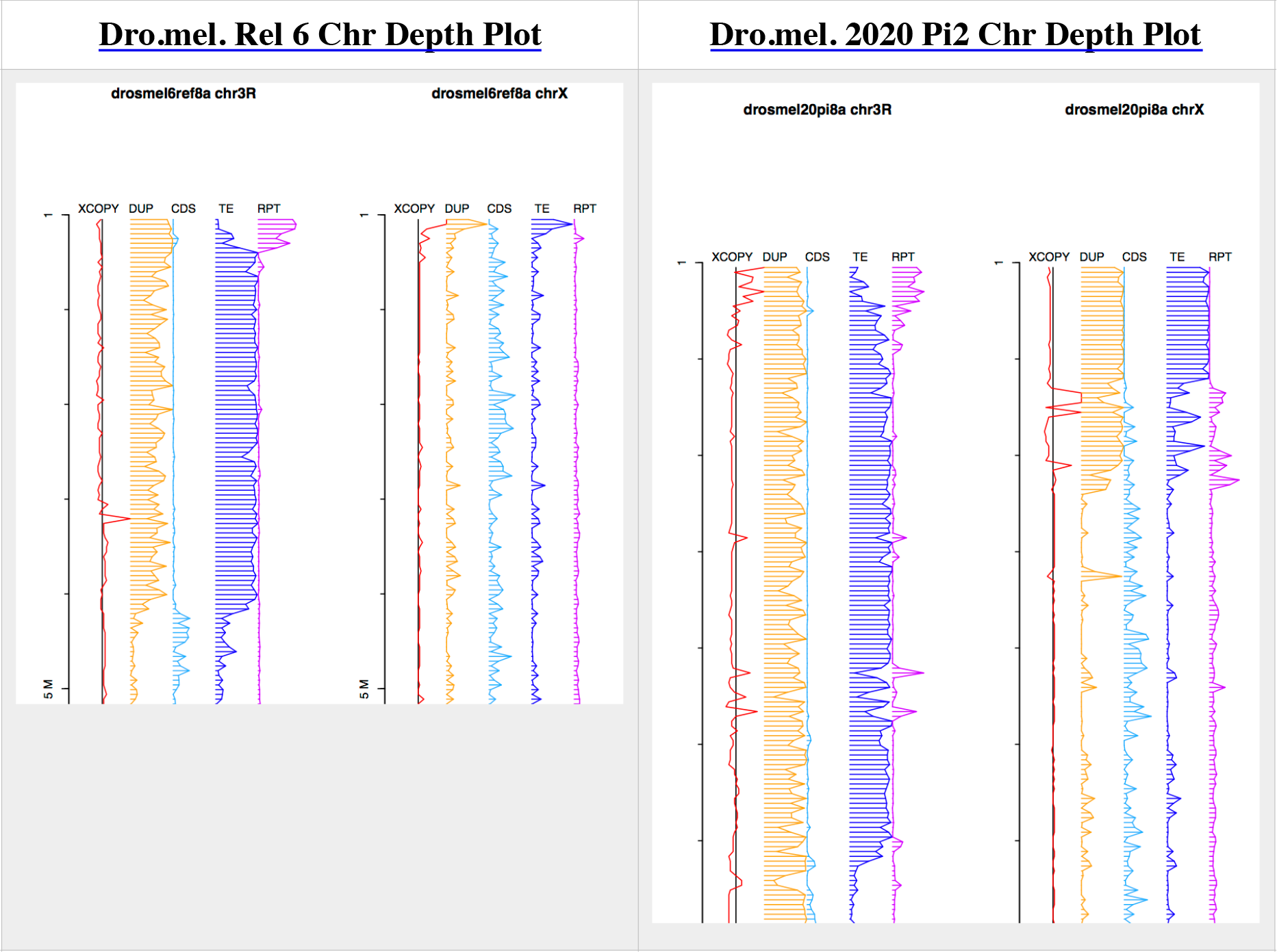
Gnodes Chromosome plot of *Drosophila melanogaster* drosmelr6, and drosmel20pi assemblies

### R4: Daphnia genome assemblies

One impetus for development of Gnodes is an aid understanding the large discrepancy in *Daphnia* water flea genome assemblies that are as small as half of flow cytometry measured sizes. Extensive gene coding sequence duplication is a likely reason that these assemblies have faltered at half-size. Half of *Daphnia* genomic DNA aligns to genes coding sequence (Figure R4.1), much more than the 10-20% of measured insects and vertebrates, or 25% in measured plants. Further details on this assessment of Daphnia genome assemblies will be reported in another paper.

**Figure R4.1.**
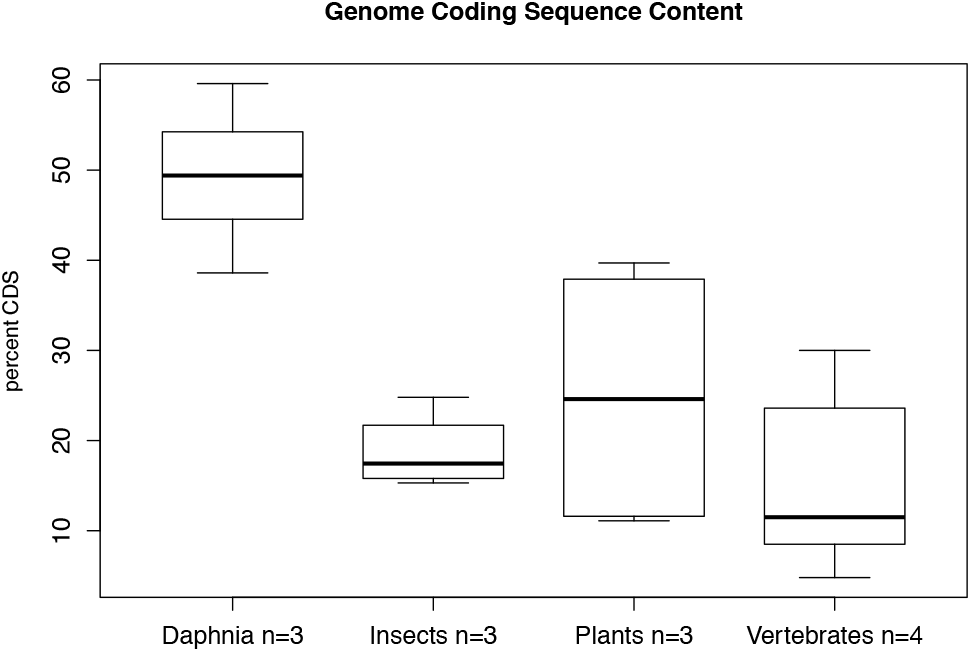
Coding sequence content of *Daphnia* waterflea genomes exceed those in sampled insects, plants and vertebrates. Box plots (median, range) of percentage of genomic DNA samples that align to coding sequences of 3 Daphnia species, and 10 other animals and plants, are plotted.

### R5: Right-sizing Daphnia genome assemblies with Gnodes

De-duplication is now a standard part of several current chromosome assemblers, often labelled as such. The rationale of de-dup is to remove heterozygous and/or excessive over-assembly sections (contigs, scaffolds), and achieve a single copy of chromosomes. EvidentialGene’s gene assembly algorithm (tr2aacds) contains the approximate equivalent of de-dup, and this author has known for over a decade [Gilbert 2013, 2019] that true gene duplications are under-represented due to this reduction (by 1%-5% depending on gene duplication level in genomes). It is clear from attempts with Daphnia DNA that chromosome assemblers suffer a similar flaw, removing true duplications, of genes, transposons and other duplicated sections, including long-read based assemblers.

This flaw with RNA assembly is not readily measurable or correctible, but with DNA assembly it is, at least measurable and to some extent correctible, where a constant depth of measured genome DNA provides the needed metric: under-assembly of true duplications have too much depth, whereas over-assembly of heterozygous or mis-assembly duplications have too little depth. This analog to counting identical twins means one looks not just at similarity but also copy depth. At least some of the de-dup algorithms lack or limit use of copy depth, in preference to simpler similarity measures.

As Gnodes measures excess and reduced copy depth over the spans of assemblies (previous sections), it may be used to pick those spans with a proper unit depth. When used with alternate assemblies, or with over-assemblies (ie, assembly before de-dup is applied), this measure combined with similarity measures allow selection of the ‘just-right’ set of contigs, and production of a chromosome assembly that nearly matches a genome experiment’s unit DNA depth. Thus Un-de-dup with Gnodes, a nearly scatological term that should be clear to those familiar with genome assembly methods, can improve genome assemblies, at both chromosome and gene levels.

A case for right-sizing by DNA depth can be seen with the fruitfly Histone gene duplication span, where the drosmel6r reference has one 100Kb span with 20-25 annotated gene copies per subunit (His1, His2a,b, His3, His4), while a newer drosmel20pi assembly expands histone duplications to 200Kb, shown in Figure R5.1. The xCopy graph (red, bottom) is above 1 for both assemblies, indicating under-assembly. Even the expanded dromel20pi is under-assembled by 2-fold in this region, according to DNA depth, which indicates 75 to 110 copies of each histone subunit in this species, rather than the 25 annotated in reference genome.

**Figure R5.1.**
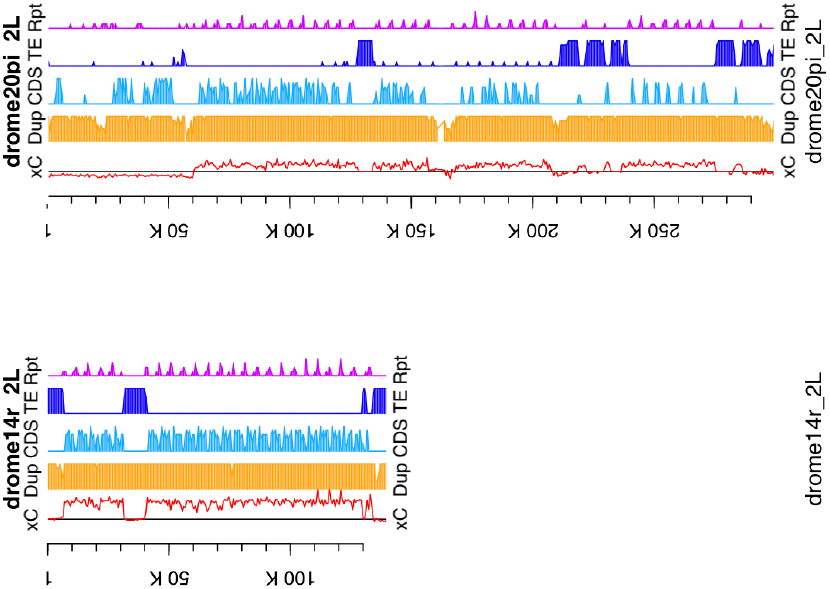
*Drosophila mel*. histone duplication span, drosmel20pi (top, 200Kb) and drosmel6r (bottom, 100 Kb) assemblies. The red graph indicates both are under-assembled, drosmel6r is 1/4, drosmel20pi is 1/2 of copy number found in DNA.

One approach to right-sizing an under-assembly like dromel14 is to replace that span with the fuller assembly span of drosmel20pi. With multiple assemblies of the same DNA sample, one can choose among fuller and under assembled contigs using DNA depth. This is the approach used to right-size a *Daphnia magna* assembly. Figure R5.2 shows the progress of re-assembling the same DNA sample of D. magna, from 1st through 6th versions, summarized by assembly and gene copy deficit plots (several more assemblies were made but discarded as not helpful). At the 4th step, an assembly made with MaSurCA provided an over-assembly, with more contigs than DNA depth warranted. This was the critical point as it allows re-selection of the contigs in this over-assembly that contain proper DNA depth, when excess duplicate contigs are removed. At the 5th step, a genome-wide average found very close approach to the right size, but inspection of details showed this to be the sum of over- and under-assembled contigs. Adjusting for this, removing more over-assembled contigs, yielded a 6th assembly that is a close approach to right-size for DNA content, though with some excess of under-assembled contigs remain. These typically are the transposon and repeat rich segments that the assemblers were not able to fully represent.

**Figure R5.2.**
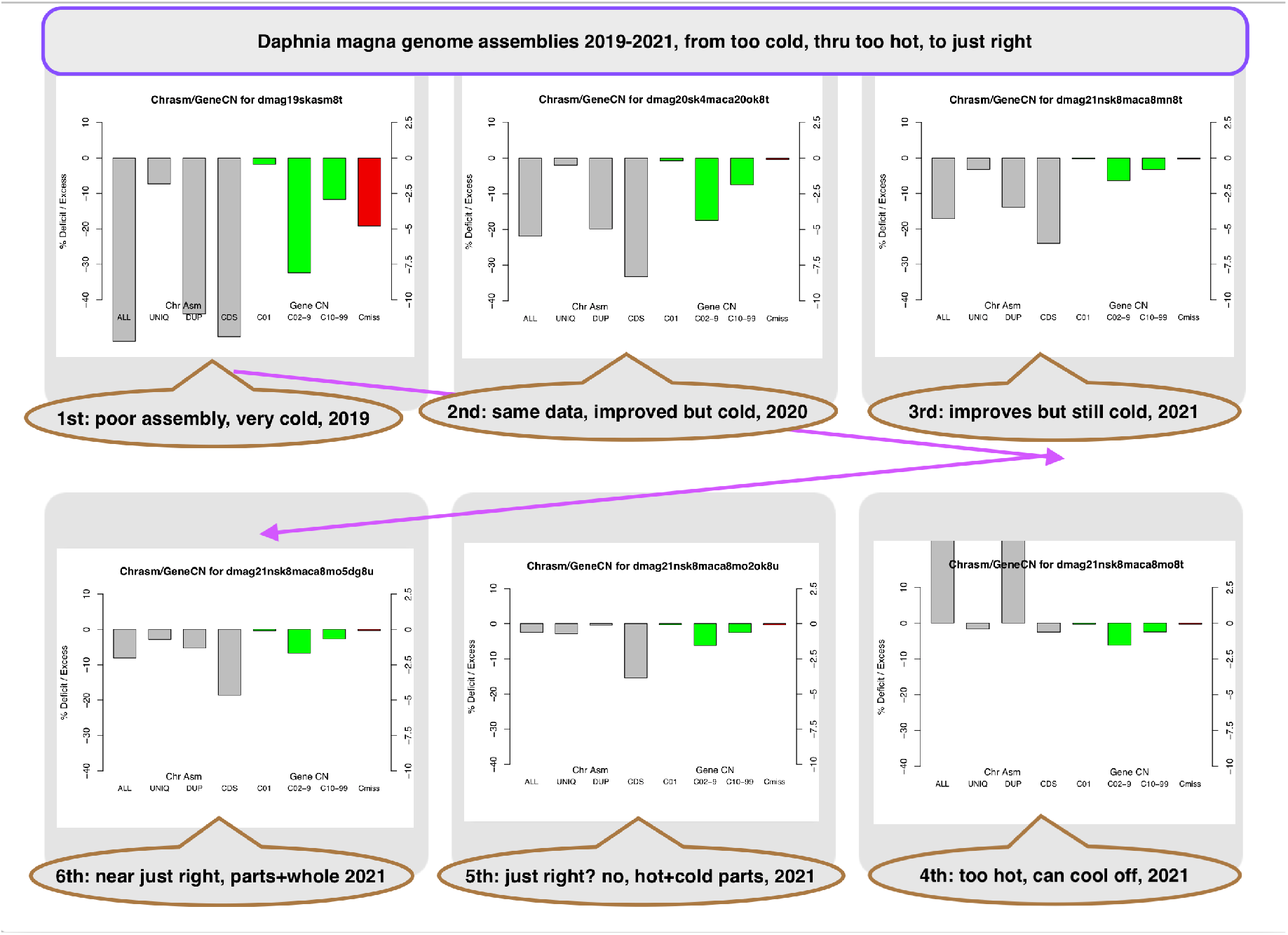
Gnodes deficit plots for *Daphnia magna* assemblies, from half-sized (110 Mb), thru over-sized, to right-sized (220 Mb).

Figure R5.3 compares the ten chromosomes, or linkage units, of *D. magna* with content depth plots of the 1st under-assembly and near-right 6th assembly, and Figure R5.4 shows details for two chromosomes. Content features similar to the 6th assembly of *D. magna* are found in sections of human, plant and fruitfly chromosomes, as found in Gnodes measurements (see Supplemental results).

### R6: Model plant, missing genome parts, found after 20 years

**Figure R5.3.**
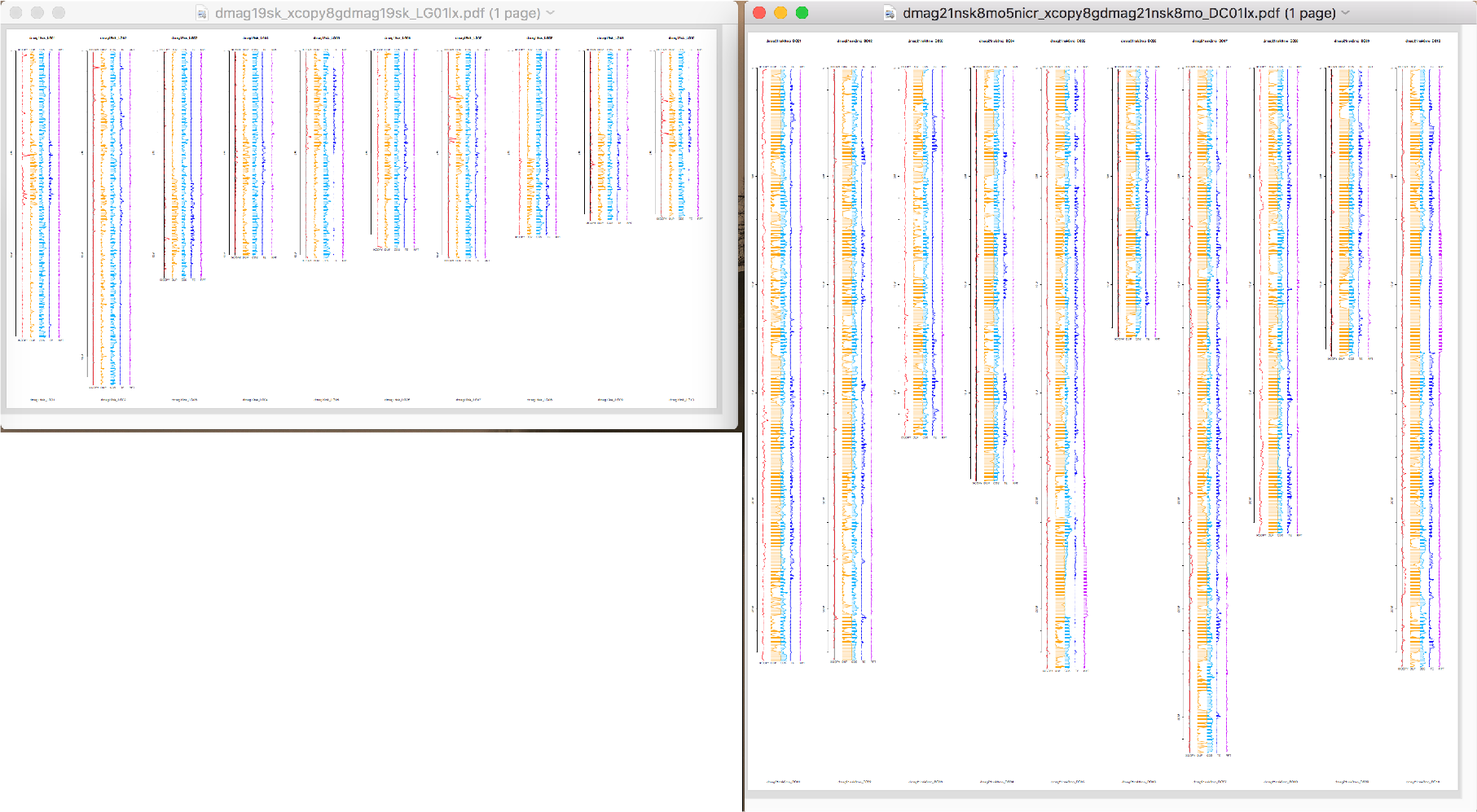
Gnodes DNA depth plots of 10 *Daphnia magna* chromosomes, 1st (110Mb) versus 6th (220 Mb) assembly, for this species with 230-390 Mb genome measured with flow cytometry. The graph columns per chromosome are Xcopy (red, left), Duplicated DNA (orange), gene CDS-mapped DNA (light blue), transposons (dark blue), and simple repeats (purple, right).

**Figure R5.4.**
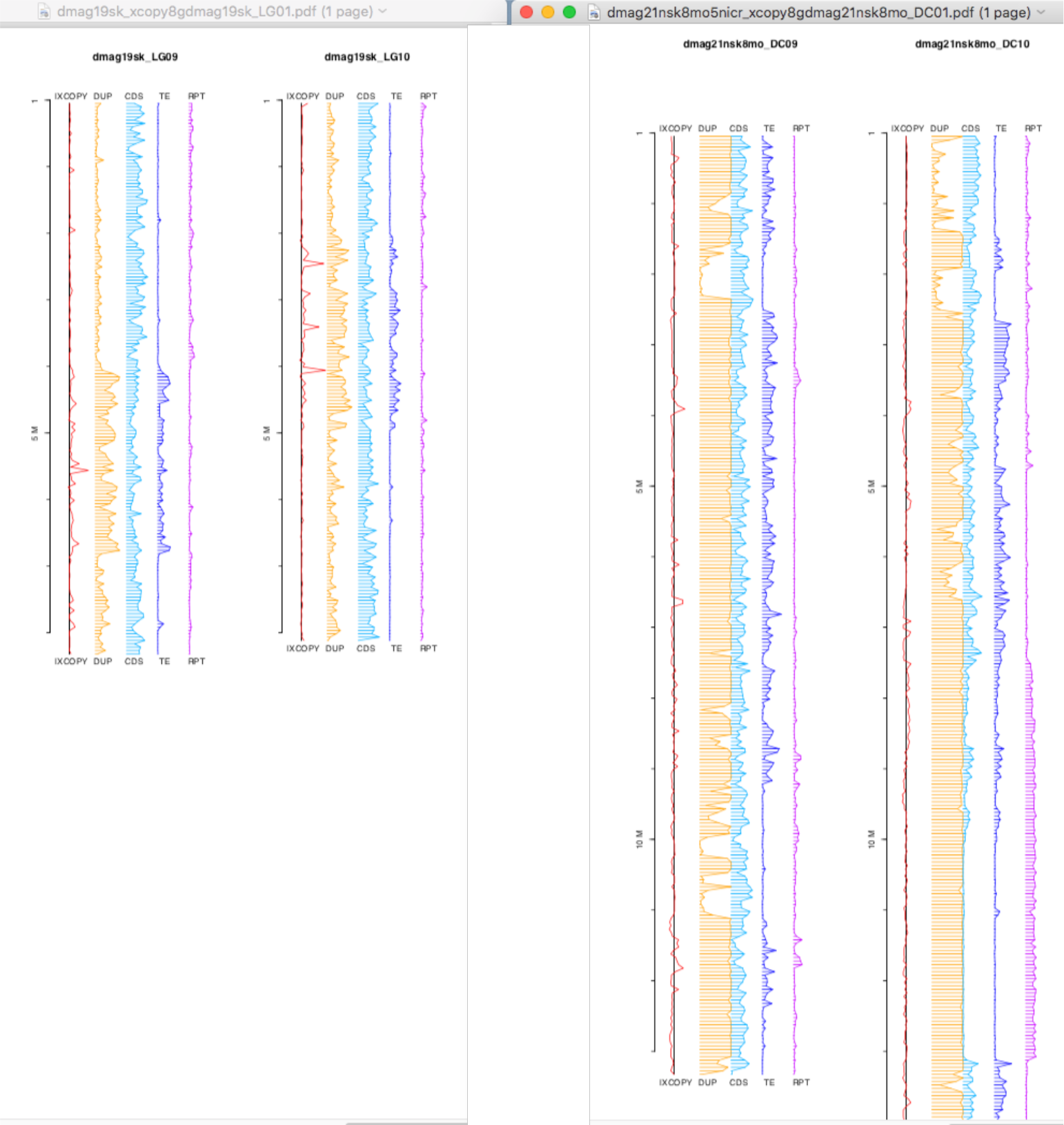
Gnodes depth detail of *Daphnia magna* chromosomes 9 and 10, 1st versus 6th assembly. Notably the 6th assembly has extensive duplicated spans (orange DUP column), larger transposon spans (dark blue, right TE). Note the appearance of large simple repeat spans (right purple RPT column) in 6th, which are common with reference species but not undersized Daphnia assemblies. Daphnia species assemblies differ from observed reference species in having extensive CDS spans (light blue middle column), unique and duplicated (percent CDS-like in genome, Figure 8).

In the year 2000, the Arabidopsis Genome Initiative published [A.G.I., 2000] a nearly complete assembly of the 5 chromosomes of this model plant. This effort entailed Sanger sequencing of BACs spanning the chromosomes, and resulted in high quality genome sequence that, while it has been improved in details, has not changed in major content nor size for 20 years. However, several of the BACs were found in 2000 to be too repetitive to sequence at the time and left undone, without apparent resolution. What and how much of the Arabidopsis genome is contained in those missing parts?

Bennett, Leitch, Price and Johnston (2003) authored a careful assessment of this genome size discrepancy, comparing flow cytometry measures with sequence assembly estimates. They found 157 Mb of genome by flow cytometry, versus 125 Mb in the assembled (119Mb) + estimated missing assembly from AGI. This discrepancy has held for 20 years; this author searched NCBI genome databases for recent At genome assemblies thru 2020, finding one a bit larger (130 Mb, At20max), but is by Gnodes analysis of poorer quality than At18tair.

Among points Bennett et al discuss:

*Was 125 Mb a significant underestimate for the arabidopsis genome? Yes, as Naish et al. [2021-Dec] add 13 Mb in centromeres (133 Mb total) but leave unfinished other missing duplicated regions.

* Is the total DNA content of the two arabidopsis NORs approx. 7.3 Mb? These are two rRNA repeat regions, at tops of Chr2, 4. Recent published data contain even larger NORs at 10 Mb, supported by re-assembly of pacbio-hifi sequence that equalizes gDNA coverage depth in these regions.

* Is the total DNA content of centromeric gaps only approx. 3 Mb? No, as the recent paper [Naish et al, 2021-dec] describes careful centromere assembly, spanning about 2.5 Mb on each of 5 chromosomes, ie 12-15 Mb total, which agrees with estimates reported by Bennett et al.

These difficulties bear on many genome projects; plant and animal genomes have a variable amount of duplicated matter that is difficult to measure and assemble, and advent of longer read sequencing has yet to resolve this in many cases (e.g. Daphnia waterfleas). A recent update to At chromosome assemblies [Naish et al., 2021-Dec] includes centromeres, one of the large, repetitive blocks missing since 2000.

Using Gnodes measures, the AGI pseudo-chromosomes (up to 2018) are found to be well resolved in the euchromatic, gene-rich, mostly unique sequence arms of all 5 chromosomes. Transposon rich spans are mostly unique sequence, and have a peak near centromeres of each chromosome. This assembly however shows huge spikes or hot-spots of un-assembled DNA copy in certain areas, where the centromeres should be, and at top telomeric ends of Chr 2 and 4, and certain other hot spots, mostly near to centromeres or ends Table R6.1 provides content summaries of three assemblies, *at18tair* known as TAIR10, 2018 update, *at21ncbi* adds centromeres and some of telomeres, using very long reads of Naish et al 2021, and *at22canu* a re-assembly of those very long reads using Canu adds Chr 2,4 telomeric rRNA, for this paper. Table R6.2 indicates the differences in chromosome parts measured in these assemblies. Table R6.3 enumerates repeat contents that differ by chromosome assembly.

**Table R6.1.**
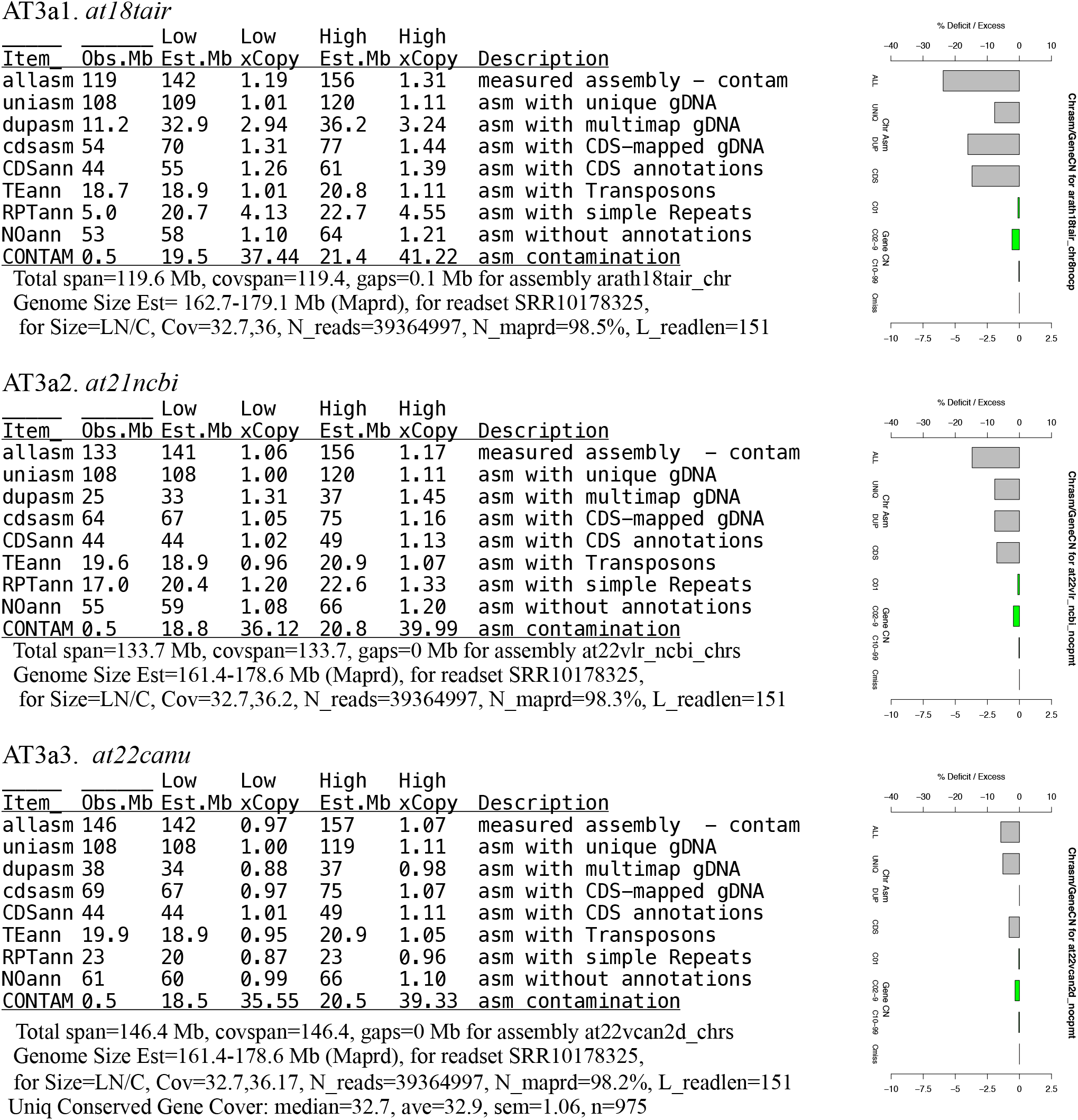
Whole genome measurement summaries for Arabidopsis assemblies, outputs of Gnodes. The assemblies are *at18tair* (AGI 2000 reference updated to 2018, AT3a1), *at21ncbi* (Naish et al 2021, AT3a2), and *at22canu* (AT3a3), described in text. Gnodes Assembly + Gene Copy Deficit plots of these tables are at right side. Low and High xCopy are based on two coverage depth measures of unique conserved genes, and bracket a likely true value; Low, High Est.Mb are estimated sizes from xCopy * Observed Mb. Items are uniqasm, dupasm from single or multimap gDNA reads, which add to allasm; TE, RPT from Repeatmasker; cdsasm, CDSann from gene-CDS mapping; NOann is part without CDS, TE, or RPT; CONTAM is chloroplast + mitochondia sequence, removed from calculation of other items. These results are for SRR10178325 read set; several others give near equivalents (Supplemental results).

**Figure R6.1.**
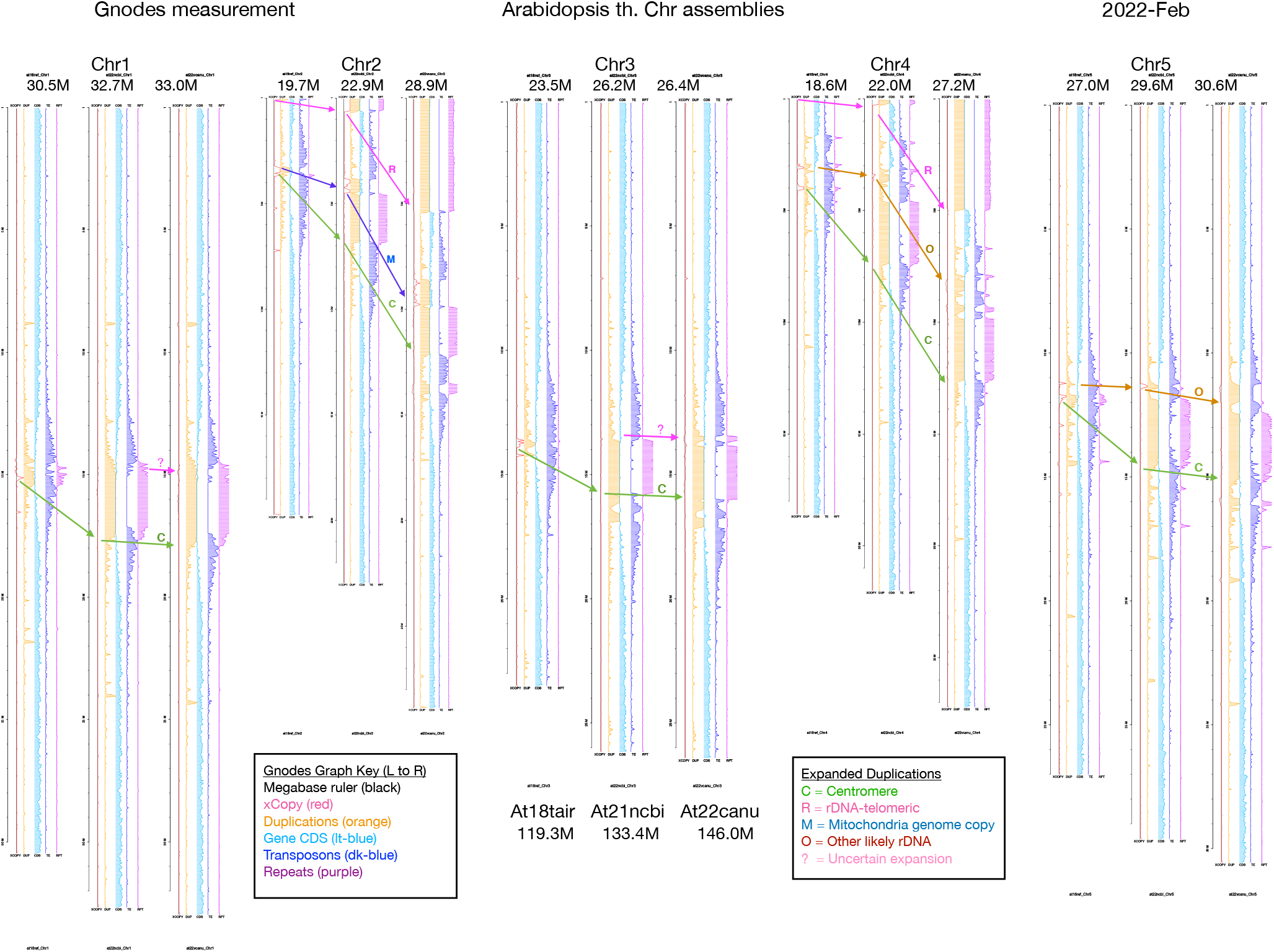
Gnodes DNA Depth graphs of 3 assemblies of Arabidopsis chromosomes 1 to 5. Regions expanded from At18tair are marked with arrows. At18tair (left) is reference (GCA_000001735.2), updated from published in 2000 thru 2018; At21ncbi (middle) is GCA_020911765.1 from Naish et al. [2021-Dec], At22canu (right) is re-assembly for this paper, available in supplement, of pacbio-hifi data of Naish et al using canu 2.2. At22canu has the most even DNA depth, reflecting assembly xCopy closest to 1 in Table R6.1 ; xCopy hot spots (>>1) are observed at expansion regions in smaller assemblies.

The large hot-spots of DNA depth (un-assembled in At18tair, expanded in other two) are indicated with arrows in Figure R6.1, where the centromeres are (C-green), and rDNA or NORs at top telomeric ends of Chrs 2, 4 (R-purple). There are certain other hot spots, mostly near to centromeres or ends. Chr2 pre-centromere has a mitochondria DNA insert: this may be one copy of 370 Kb mtDNA (at21ncbi), or multiple copies (at22canu). Also, at22canu contains two other pre-centromere expansions of 500-700 Kb (O, brown), on Chr4 and Chr5, apparently rDNA repeats. These are near identical to At 5S rDNA cluster (AB073495) and sRNA locus (LS474512). There are also some uncertain, smaller expansions in this at22canu draft assembly.

**Table R6.2.** Chromosome parts measures of At. assemblies, in megabases, plotted in Figure R6.1. The at00agi rows are those produced by initial assembly of A.G.I., 2000, including observed and estimated sizes from that report. Estimated sizes for others come from Gnodes, for at18tair of TAIR10 updated in 2018, at21ncbi and at22canu described in text.

| ChrID | ObsLoc | EstLoc | Nuniq | Ndup | Ncds | NCDSan | NTEan | NRPTan | NNOan |
| --- | --- | --- | --- | --- | --- | --- | --- | --- | --- |
| at00agi_Chr1 | 30.4 | 35.0 |  |  |  |  |  |  |  |
| at18tair_Chr1 | 30.5 | 34.1 | 27.3 | 2.9 | 14.3 | 11.6 | 3.7 | 1.4 | 14.0 |
| at21ncbi_Chr1 | 32.7 | 36.0 | 27.4 | 5.3 | 16.3 | 11.6 | 3.7 | 3.8 | 14.0 |
| at22canu_Chr1 | 33.0 | 36.0 | 27.4 | 5.5 | 16.5 | 11.6 | 3.7 | 4.0 | 14.1 |
| at00agi_Chr2 | 19.7 | 22.0 |  |  |  |  |  |  |  |
| at18tair_Chr2 | 19.7 | 29.7 | 17.8 | 1.9 | 8.1 | 6.6 | 3.9 | 0.7 | 8.8 |
| at21ncbi_Chr2 | 22.9 | 31.8 | 17.7 | 5.2 | 10.3 | 6.6 | 4.0 | 3.2 | 9.4 |
| at22canu_Chr2 | 28.9 | 30.3 | 17.6 | 11.2 | 12.9 | 6.7 | 4.1 | 6.3 | 12.0 |
| at00agi_Chr3 | 23.5 | 25.5 |  |  |  |  |  |  |  |
| at18tair_Chr3 | 23.5 | 34.7 | 21.3 | 2.1 | 10.6 | 8.6 | 4.0 | 0.8 | 10.4 |
| at21ncbi_Chr3 | 26.2 | 28.8 | 21.3 | 4.8 | 12.4 | 8.6 | 4.1 | 2.9 | 10.9 |
| at22canu_Chr3 | 26.4 | 28.8 | 21.3 | 5.1 | 12.6 | 8.6 | 4.1 | 3.1 | 10.9 |
| at00agi_Chr4 | 18.6 | 20.8 |  |  |  |  |  |  |  |
| at18tair_Chr4 | 18.6 | 24.7 | 16.8 | 1.8 | 8.3 | 6.7 | 3.3 | 0.8 | 8.0 |
| at21ncbi_Chr4 | 22.0 | 26.4 | 16.8 | 5.1 | 10.8 | 6.7 | 3.6 | 3.6 | 8.3 |
| at22canu_Chr4 | 27.2 | 28.1 | 16.8 | 10.4 | 12.8 | 6.7 | 3.6 | 6.4 | 10.8 |
| at00agi_Chr5 | 27.0 | 31.3 |  |  |  |  |  |  |  |
| at18tair_Chr5 | 27.0 | 32.8 | 24.6 | 2.4 | 12.5 | 10.2 | 3.9 | 1.2 | 12.1 |
| at21ncbi_Chr5 | 29.6 | 33.1 | 24.6 | 5.0 | 14.3 | 10.2 | 4.2 | 3.4 | 12.2 |
| at22canu_Chr5 | 30.6 | 33.2 | 24.6 | 5.9 | 14.5 | 10.2 | 4.4 | 3.5 | 12.8 |
| ChrID | ObsLoc | EstLoc | Nuniq | Ndup | Ncds | NCDSan | NTEan | NRPTan | NNOan |
| at00agi_Total | 119.1 | 134.6 |  |  |  |  |  |  |  |
| at18tair_Total | 119.3 | 155.9 | 107.8 | 11.2 | 53.8 | 43.7 | 18.7 | 5.0 | 53.3 |
| at21ncbi_Total | 133.4 | 156.1 | 107.8 | 25.4 | 64.1 | 43.6 | 19.6 | 17.0 | 54.8 |
| at22canu_Total | 146.0 | 156.4 | 107.7 | 38.2 | 69.4 | 43.8 | 19.9 | 23.4 | 60.6 |

Column key: ObsLoc = observed size, EstLoc = estimated size, Nuniq, Ndup, Ncds: assembly with unique, multimapped and CDS-mapped gDNA (as per whole genome summaries), NCDSan, NTEan, NRPTan, NNOan : assembly with annotations of gene CDS, Transposons, simple Repeats, and no annotations.
+ Cols w/ diff: Ndup, Ncdsasm (but not NCDSann), NRPTan, NNOan (some rDNA here?)
- Cols w/ same: Nuniq (but small Chr2), NCDSan, NTEan
at00agi statistics are at https://www.arabidopsis.org/portals/genAnnotation/gene_structural_annotation/agicomplete.jsp

**Table R6.3.** Repeats differing by chromosome assembly, from RepeatMasker. Retroelements and Transposons, Low_complexity, Simple_repeat, snRNA, tRNA, Other all have nearly same values. Supplement has full RepeatMasker tables.

| Chr1 | Chr2 | Chr3 | Chr4 | Chr5 | Type | Assembly |
| --- | --- | --- | --- | --- | --- | --- |
| 1077 | 206 | 104 | 1236 | 919 | Satellite | at18tair |
| 9 | 4 | 4 | 2 | 3 | rRNA | at18tair |
| 22006 | 17179 | 14527 | 18247 | 10417 | Satellite | at21ncbi *** |
| 7 | 124 | 1 | 68 | 3 | rRNA | at21ncbi |
| 24317 | 20598 | 15960 | 19280 | 10777 | Satellite | at22canu *** |
| 7 | 1202 | 1 | 1039 | 3 | rRNA | at22canu *** |

### R7: Genome size estimations, redux

The most accurate estimator of animal and plant genome sizes from DNA sequence samples, as measured by agreement with flow cytometry estimates, is the simple formula **G = L*N/*c*** (Lander and Waterman, 1988, as ***c* = L*N/G**), where ***c*** is an accurate measure of a presumed constant DNA depth over nuclear chromosomes, and L*N (read length times number of reads, or total bases sampled), adheres to assumptions of even depth of coverage, no significant contaminants nor sequencing errors.

**Table R7.1.** Genome size estimations.

| Genome_Name | SRA_Reads | Genome Size Estimations, % of FCsize |  |  |  |  |  |  |  |
| --- | --- | --- | --- | --- | --- | --- | --- | --- | --- |
|  |  | FCsize | LNCm | LNCa | ASM | GScop | fiGSE | MGSE | covest |
| Arath18TAIR | SRR10178325 | 157 | 96 | 97 | 76 | 91 | 92 | 87 | 96 |
| DrosMel14 | SRR11460802 | 176 | 93 | 96 | 81 | 84 | 87 | na | 90 |
| Daphplx19ml | SRR1333379n | 200 | 98 | 104 | 92 | 76 | 0 | na | 168 |
| Daphmag20skm | SRR7825549 | 234 | 97 | 106 | 95 | 76 | 98 | 76 | 146 |
| Cucumber19CGI | SRR11300859 | 372 | 95 | 99 | 61 | 69 | 81 | 80 | 145 |
| Beet18Usda | SRR6305245 | 705 | 99 | 101 | 77 | 73 | 80 | 87 | na |
| Chicken19GRC | SRR3954707 | 1223 | 93 | 94 | 86 | 82 | 89 | na | 92 |
| Corn20MGD5 | ERR3288215 | 2352 | 98 | 102 | 93 | 64 | 70 | 47 | na |
| Pig11ref | SRR4341337 | 2924 | 100 | 105 | 85 | 78 | 95 | na | 102 |
| Human19GRC | SRR12898282 | 3423 | 95 | 97 | 88 | 77 | 81 | 88 | 146 |
|  | Mean | 1176 | 96.4 | 100.1 | 83.4 | 77.0 | 85.9 | 77.5 | 123.1 |
|  | StDev | 1257 | 2.4 | 4.1 | 10.2 | 7.6 | 8.8 | 15.7 | 31.1 |

| Genome_Name | Genome Measurements by Gnodes |  |  |  |  |  |  |
| --- | --- | --- | --- | --- | --- | --- | --- |
|  | Cucg | Dupasm | CDSasm | TEann | RPTann | Map% | tabdate |
| Arath18TAIR | 34.3 | 23 | 50 | 13 | 15 | 98.5 | 2022_Jan_26 |
| Fruitfly14 | 33.1 | 23 | 25 | 16 | 12 | 96.7 | 2022_Jan_26 |
| Daphnia_p19l | 42.3 | 52 | 60 | 6 | 7 | 94.5 | 2021_Oct_3 |
| Daphnia_m20m | 48.7 | 61 | 76 | 12 | 4 | 91.7 | 2021_Sep_24 |
| Cucumber19 | 23.9 | 48 | 33 | 3 | 9 | 96.7 | 2022_Feb_15 |
| Beet18USDA | 131. | 63 | 17 | 7 | 5 | 98.4 | 2022_Mar_5 |
| Chicken19GRC | 27.9 | 10 | 8 | 10 | 7 | 98.9 | 2021_Oct_7 |
| Corn20MGDB | 18.1 | 70 | 16 | 73 | 4 | 96.9 | 2022_Mar_6 |
| Pig11Ref | 19.3 | 15 | 27 | 42 | 9 | 94.8 | 2021_Oct_8 |
| Human19GRC | 20.1 | 15 | 14 | 51 | 7 | 98.2 | 2021_Oct_20 |
| Mean | 39.9 | 38.0 | 32.6 | 23.3 | 7.9 | 96.5 |  |
| StDev | 33.6 | 23.0 | 22.3 | 23.6 | 3.5 | 7.9 |  |

LNCall, measured with all DNA bases including those unmapped to assembly, would be the best estimator of FCsize, when unmapped DNA is mostly organismal, not contaminant or artifact. See Maprate vs LNCall, LNCmap. Many genome DNA samples have contaminants in large amounts, including Daphnia with bacteria, and plants with 1000s of chloroplast genomes/sample. Gnodes measures of plant DNA were decontaminated by mapping to chloroplast assemblies.

The deviation of K-mer estimates from FCsize is large, with a wide range of estimates. On average these are under-estimates, along with under-assembled chromosome assemblies. covest, modeling for repetitive DNA as run here, produced over-estimates. FindGSE among K-mer methods has the closest approach to FC measured sizes, but with a large range in estimates, only 3/10 are within 10% of measured size. MGSE, the other map-based estimator here [Pucker 2019], has a larger error than Gnodes LNC, due to its simpler but less precise method of measuring Cucg, the constant depth of DNA at unique genes. DNA duplications are suspected as a primary problem for the K-mer methods, but the associated measures produced by Gnodes do not support that alone (Dupasm) as the problem, and neither amounts of transposons nor simple repeats correspond with variance in K-mer estimates.

### R8: Not all genome DNA contains a genome

There is good agreement between FC and Gnodes when DNA samples are those used to assemble chromosomes, as in Table R7.1. Not all DNA samples have such agreement. The main reason? Not all DNA samples have a complete genome sample: sampling molecular methods can degrade or skew the genomic content, e.g. PCR amplification; and animal, plant bio-samples from studies that manipulate genomes, e.g. mutations, inbred and hybrid lines, do change genomes, sometimes in large ways. Table R8.1 presents some details relating to DNA samples used for measurement of genomes.

**Table R8.1.**
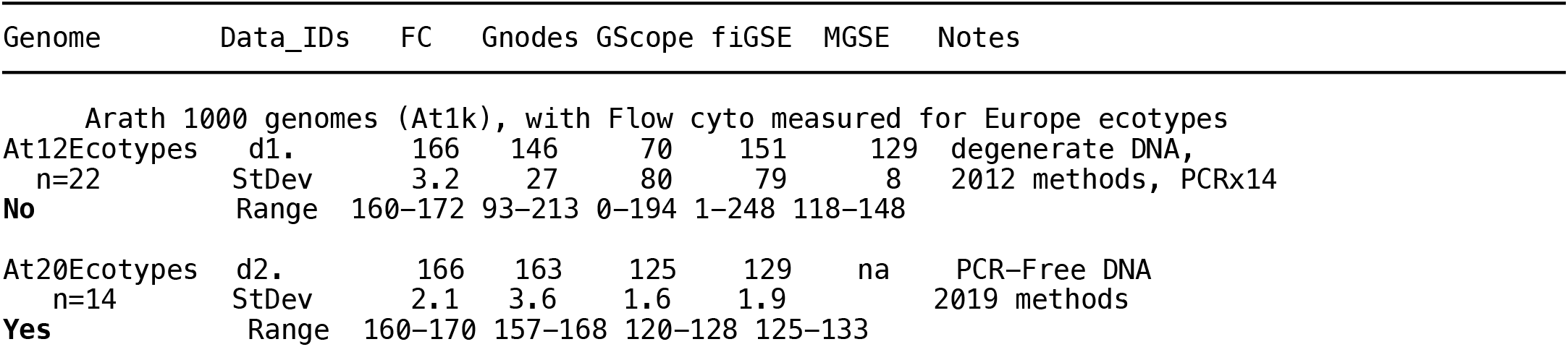
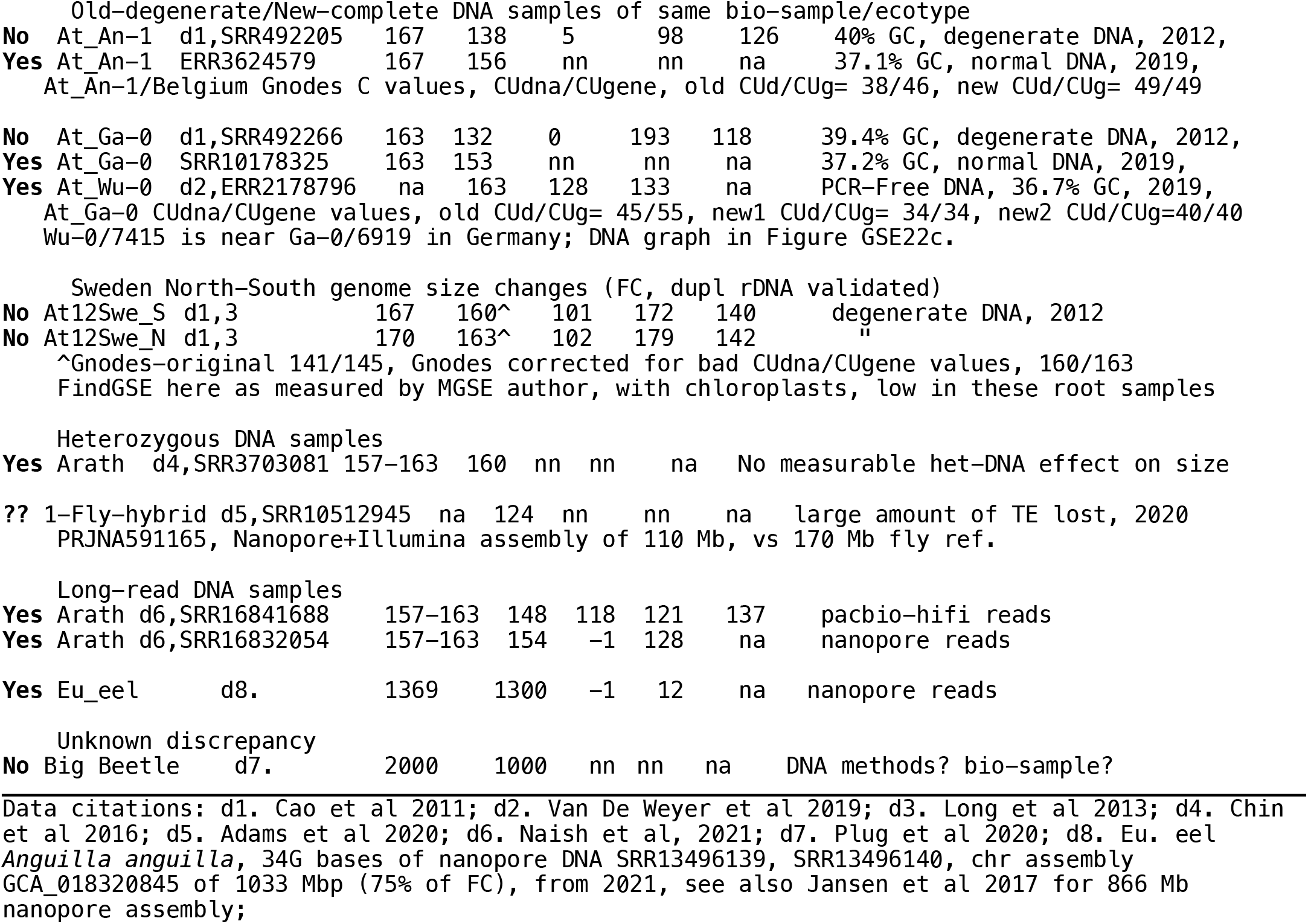
Genome DNA samples that do or don’t contain full genomes, with flow cytometry (FC) sizes, Gnodes LNCall, GenomeScope (GScope), findGSE (fiGSE) and MGSE size estimates. Yes/No indicates if the DNA samples have full genomes.

| Genome | Data_IDs | FC | Gnodes | GScope | fiGSE | MGSE | Notes |
| --- | --- | --- | --- | --- | --- | --- | --- |
| Arath 1000 genomes (At1k), with Flow cyto measured for Europe ecotypes |  |  |  |  |  |  |  |
| At12Ecotypes | d1. | 166 | 146 | 70 | 151 | 129 | degenerate DNA, |
| n=22 | StDev | 3.2 | 27 | 80 | 79 | 8 | 2012 methods, PCRx14 |
| No | Range | 160-172 | 93-213 | 0-194 | 1-248 | 118-148 |  |
| At20Ecotypes | d2. | 166 | 163 | 125 | 129 | na | PCR-Free DNA |
| n=14 | StDev | 2.1 | 3.6 | 1.6 | 1.9 |  | 2019 methods |
| Yes | Range | 160-170 | 157-168 | 120-128 | 125-133 |  |  |

Old-degenerate/New-complete DNA samples of same bio-sample/ecotype
|  |  |  |  |  |  |  |  |  |
| --- | --- | --- | --- | --- | --- | --- | --- | --- |
| No | At_An-1 | d1,SRR492205 | 167 | 138 | 5 | 98 | 126 | 40% GC, degenerate DNA, 2012, |
| Yes | At_An-1 | ERR3624579 | 167 | 156 | nn | nn | na | 37.1% GC, normal DNA, 2019, |
|  | At_An-1/Belgium Gnodes C values, CUDna/CUgene, old CUD/CUG= 38/46, new CUD/CUG= 49/49 |  |  |  |  |  |  |  |
| No | At_Ga-0 | d1,SRR492266 | 163 | 132 | 0 | 193 | 118 | 39.4% GC, degenerate DNA, 2012, |
| Yes | At_Ga-0 | SRR10178325 | 163 | 153 | nn | nn | na | 37.2% GC, normal DNA, 2019, |
| Yes | At_Wu-0 | d2,ERR2178796 | na | 163 | 128 | 133 | na | PCR-Free DNA, 36.7% GC, 2019, |
|  | At_Ga-0 CUDna/CUgene values, old CUD/CUG= 45/55, new1 CUD/CUG= 34/34, new2 CUD/CUG=40/40 |  |  |  |  |  |  |  |
|  | Wu-0/7415 is near Ga-0/6919 in Germany; DNA graph in Figure GSE22c. |  |  |  |  |  |  |  |

Sweden North-South genome size changes (FC, dupl rDNA validated)
|  |  |  |  |  |  |  |  |  |
| --- | --- | --- | --- | --- | --- | --- | --- | --- |
| No | At12Swe_S | d1,3 | 167 | 160^ | 101 | 172 | 140 | degenerate DNA, 2012 |
| No | At12Swe_N | d1,3 | 170 | 163^ | 102 | 179 | 142 | " |

^Gnodes-original 141/145, Gnodes corrected for bad CUDna/CUgene values, 160/163 FindGSE here as measured by MGSE author, with chloroplasts, low in these root samples

Heterozygous DNA samples
|  |  |  |  |  |  |  |  |  |
| --- | --- | --- | --- | --- | --- | --- | --- | --- |
| Yes | Arath | d4,SRR3703081 | 157-163 | 160 | nn | nn | na | No measurable het-DNA effect on size |

|  |  |  |  |  |  |  |  |  |
| --- | --- | --- | --- | --- | --- | --- | --- | --- |
| Yes | Arath | d6,SRR16841688 | 157-163 | 148 | 118 | 121 | 137 | pacbio-hifi reads |
| Yes | Arath | d6,SRR16832054 | 157-163 | 154 | -1 | 128 | na | nanopore reads |
| Yes | Eu_eel | d8. | 1369 | 1300 | -1 | 12 | na | nanopore reads |

Unknown discrepancy
|  |  |  |  |  |  |  |  |  |
| --- | --- | --- | --- | --- | --- | --- | --- | --- |
| No | Big Beetle | d7. | 2000 | 1000 | nn | nn | na | DNA methods? bio-sample? |

Data citations: d1. Cao et al 2011; d2. Van De Weyer et al 2019; d3. Long et al 2013; d4. Chin et al 2016; d5. Adams et al 2020; d6. Naish et al, 2021; d7. Plug et al 2020; d8. Eu. eel *Anguilla anguilla*, 34G bases of nanopore DNA SRR13496139, SRR13496140, chr assembly GCA\_018320845 of 1033 Mbp (75% of FC), from 2021, see also Jansen et al 2017 for 866 Mb nanopore assembly;

The Arabidopsis 1000 genomes project (At1k) included a set of ecotype population samples with flow cytometry measures [Cao et al 2011; Long et al 2013]. These have been used by authors of findGSE and MGSE to validate their methods, and offered a like sample for Gnodes. Gnodes turned up a large discrepancy of very large ranges in size, and average size well below other Arath DNA samples. Details of this discrepancy turned up a key indicator: C coverage depth values for “unique DNA” (CUdna) are well below C coverage for unique conserved genes (CUgene or Cucg). This contrasts to other DNA samples and a basic premise of CUdna >= CUgene. That is, measured unique spans of genome assemblies should contain the same DNA depth as unique genes, or greater depth when they have un-assembled duplications. A much lower coverage in non-genic regions is a puzzle. For the well-assembled euchromatic, gene-bearing spans of Arath, this can’t be explained by mis-assembly, nor can the ecotype population differences in this species explain such a very large loss of non-genic DNA.

More recent DNA samples of Arath ecotypes obtained with PCR-Free methods [Van de Weyer et al 2019], similar but not identical with the 2012 At1k set with flow cytometry, were then examined, and the discrepancy of CUdna << CUgene disappeared.

Figure R8.1 presents a detail comparison of such bad DNA (missing AT rich, gene-poor sections) for one of several old Arath samples with flow cytometry data that has been used in other genome size estimation studies. The CDS and GC-content ups and downs coincide with peaks and troughs in old DNA sample. Newer DNA samples have an expected near-1 coverage depth ratio over all these mostly-unique spans. Statistics for these two spans of 200Kb on Chr3, with CDS versus no CDS, are GC% 39.6%+/-0.29 vs 31.9%+/-0.23, new DNA xCopy 1.0+/-0.01 vs 1.0+/-0.01, old DNA xCopy 1.0+/-0.01 vs 0.51+/-0.01. Thus old DNA averages 1/2 of expected coverage in non-CDS spans. This indicated a problem with AT-rich, non-genic regions such as that reported in [Ji et al 2014]. That study found PCR-amplification of several cycles produced a significant under–representation of AT-rich DNA. PCR-amplification of 14x was used with old At 2012 DNA. PCR-Free samples for the Arath ecotypes from 2019 do not show this loss of AT-rich regions.

**Figure R8.1.**
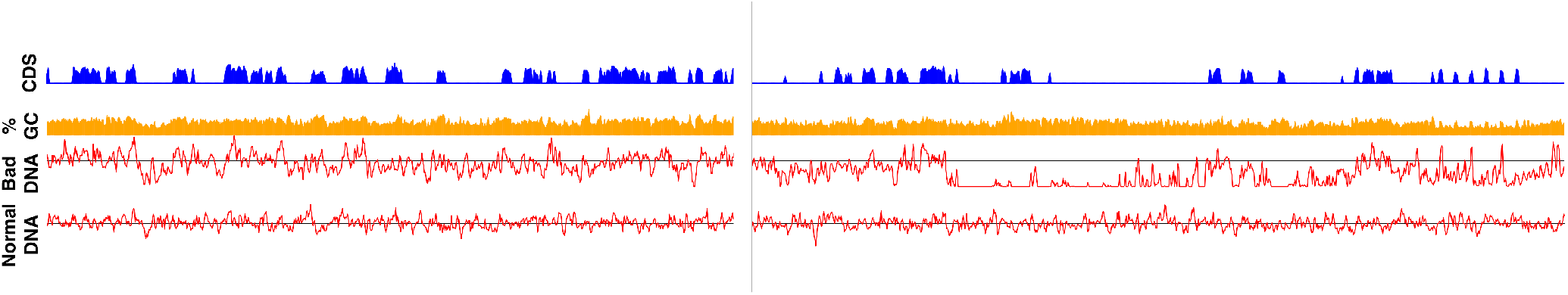
Example coverage graphs for 2012 versus 2019 DNA samples of same bio-sample (At_GA-0) at two euchromatic spans on *Arabidopis thal*. Chr3, regions that are well-assembled and gene rich. Old 2012 is middle red graph, “bad DNA” of SRR492266, using PCR of 2012. New 2019 is bottom red graph “normal DNA” of SRR10178325. Black bar in these graphs is the 1.0 coverage value of unique conserved genes of these samples; graphs below that indicate a deficit in DNA content. GC content, orange percentage, and coding sequence content (blue CDS) are displayed above; their ups and downs coincide with peaks and troughs in old DNA sample. New DNA sample has an expected near-1 coverage depth ratio over all these mostly-unique spans.

One outcome from this is that those At1k samples apparently useful for comparing GSE methods with flow cytometry are no good, as the DNA samples are incomplete. However among those At1k samples a clear case of population genome size shifts was reported [Long et al 2013] for North versus South Sweden populations. The flow cytometry differences coincide with measured changes in the large duplications of ribosomal DNA (esp. 45s rDNA on tips of chromosomes 2,4) amounting to a 3 Mb average N-S divide. Gnodes also measures that 3 Mb difference, though due to degenerate DNA samples the overall sizes are lower than true values. A computational correction for Gnodes is possible in this case, based on the CUdna/CUgene discrepancy, which recovers the measured FC genome sizes and average 3 Mb difference (Table R8.1, N-S values for FC are 170-167 Mb, for Gnodes-corrected 163-160 Mb). Due to large variance, N-S difference for Gnodes and MGSE are not statistically significant, but do have significant correlations with FC sizes for this data set. The K-mer estimates do not have significant correlations nor size differences, though findGSE has the direction of change.

The example of European eel (*Anguilla anguilla*) is useful: It has a flow cytometry + other molecular measures averaging 1369 Mb. An early use of nanopore reads [Jansen et al 2017] assembled 860 Mb (66% of FC), with GenomeScope estimates near that. More recent Nanopore reads assembled to 1033 Mbp (75% of FC), and the Gnodes estimate measured from this long-read set matches the FC estimate, as does size estimated from older Illumina reads. The K-mer estimators fail with this long-read data, which has a unique gene coverage depth of 25. Discrepancies in genome parts between new assembly and Gnodes measures is small, and spread over unique and duplicated spans, CDS, transposons and repeats, suggesting this new assembly is closer to accurate.

The case of one fruitfly with a smaller genome than reference genome is interesting; it is a hybrid of two inbred lines, sequenced and assembled recently [Adams et al 2020]. The authors find that most major genome portions are present in their assembly, but for large amounts of transposon regions, and speculate that transposons caused assembly problems. However Gnodes measures that their DNA sample of this one fly lacks this transposon DNA, so one can instead speculate there was a biological loss, perhaps large deletions of incompatible parental segments. In *Drosophila melanogaster*, heterozygous deletions are well-tolerated [Cook et al 2012] and deletions of transposon-rich telomeres occur at a high rate [McGurk et al 2021].

### R9: Human genome assembly, complete or not?

In the year 2004, the complete human genome sequence was published [International Human Genome Sequencing Consortium, 2004]. In this year 2022, a more complete human genome sequence is published [Human T2T Consortium, Nurk et al., 2022]. Is this truly a complete human chromosome assembly?Likely yes, with some uncertainty, as DNA samples indicate some discrepancies.

Gnodes analyses (Table R9.1) indicate up to 5% DNA contents may not be accounted for (high xCopy =1.05 of Hs22c), however the reasonable low estimate is in perfect agreement with the T2T assembly of 3117 Mb (low xCopy = 1.00 of Hs22c). Genes copy number analysis of Gnodes identifies 70 duplicated genes in the T2T DNA sample that are under-represented in the assembly, including proteins of beta-defensin, chromosomal, glyceraldehyde, heterogeneous ribonucleoproteins, high mobility group, homeobox, histone, keratin, metallothionein, myosin, olfactory receptor, peptidyl-prolyl isomerase, syncytin, and uncharacterized and zinc finger genes.

These analyses are also in agreement with flow cytometry estimates of 3423 Mb [Dolez & Greilhuber 2010]. The DNA sample is female, with a low-end GSE of 3368 for mapped reads (Table R9.1 Hs22c); corrected for missed Y-DNA, minus mitochondrial DNA, gives a low-end GSE of 3428 (G= L*N/Cm = 250* (801.694005 + 14.202451)/59.5). Unmapped reads, if human nuclear DNA, increase this to 3504. The other DNA sample used in Table R9.1 produces similar estimated sizes for both T2T and older GRC assemblies. The T2T assembly fills large centromeric and telomeric gaps of the prior assembly. Gnodes suggest components that may be a bit under-assembled include transposon spans (50 Mb), some duplicated genes, and regions unannotated in this analysis that may include non-coding genes (90 Mb).

Nurk et al. (2022) provide detailed DNA coverage analyses for their assembly with their data, analogous to Gnodes analyses, that well supports their calling this a complete genome. The analysis here is less extensive than for the T2T publication, and the discrepancies indicated can be compared and resolved.There are enough uncertainties in these measures and method details that further effort may improve this chromosome assembly. Measures provided by Gnodes may help resolve discrepancies and improve human, other animal and plant genome reconstructions.

**Table R9.1.** Human chromosome assemblies with whole genome measurement summaries of Gnodes. The assemblies are human19grc (a. GRC 2019), human22ttot (b, c. T2T Consortium, 2022), with two DNA samples SRR12898282 (a, b. female HG004) and SRR3189741,2 (c. CHM13 biosample of human22ttot). Table contents are as in Table R1.2 and Table R6.1 for plant assembly comparisons.

Hs22a. human19grc x SRR12898282 DNA of HG004
|  |  | Low | Low | High | High |  |
| --- | --- | --- | --- | --- | --- | --- |
| Item | Obs.Mb | Est.Mb | xCopy | Est.Mb | xCopy | Description |
| allasm | 2920 | 3065 | 1.05 | 3352 | 1.15 | measured assembly |
| uniasm | 2547 | 2609 | 1.02 | 2853 | 1.12 | asm with unique gDNA |
| dupasm | 374 | 456 | 1.22 | 499 | 1.34 | asm with multimap gDNA |
| cdsasm | 463 | 419 | 0.90 | 458 | 0.99 | asm with CDS-mapped gDNA |
| CDSann | 51 | 49 | 0.98 | 54 | 1.07 | asm with CDS annotations |
| TEann | 1559 | 1565 | 1.00 | 1712 | 1.10 | asm with Transposons |
| RPTann | 154 | 213 | 1.39 | 233 | 1.52 | asm with simple Repeats |
| NOann | 1178 | 1256 | 1.07 | 1374 | 1.17 | asm without annotations |

Hs22b. human22ttot x SRR12898282 DNA of HG004
|  |  | Low | Low | High | High |  |
| --- | --- | --- | --- | --- | --- | --- |
| Item | Obs.Mb | Est.Mb | xCopy | Est.Mb | xCopy | Description |
| allasm | 3069 | 3033 | 0.99 | 3398 | 1.11 | measured assembly |
| uniasm | 2559 | 2582 | 1.01 | 2893 | 1.13 | asm with unique gDNA |
| dupasm | 510 | 451 | 0.88 | 505 | 0.99 | asm with multimap gDNA |
| cdsasm | 307 | 264 | 0.86 | 295 | 0.96 | asm with CDS-mapped gDNA |
| CDSann | 51 | 48 | 0.94 | 54 | 1.05 | asm with CDS annotations |
| TEann | 1477 | 1450 | 0.98 | 1624 | 1.10 | asm with Transposons |
| RPTann | 280 | 240 | 0.86 | 269 | 0.96 | asm with simple Repeats |
| NOann | 1284 | 1315 | 1.02 | 1473 | 1.15 | asm without annotations |

Hs22c. human22ttot x SRR3189741,2 DNA of CHM13
|  |  | Low | Low | High | High |  |
| --- | --- | --- | --- | --- | --- | --- |
| Item | Obs.Mb | Est.Mb | xCopy | Est.Mb | xCopy | Description |
| allasm | 3068 | 3065 | 1.00 | 3228 | 1.05 | measured assembly |
| uniasm | 2669 | 2680 | 1.00 | 2823 | 1.06 | asm with unique gDNA |
| dupasm | 399 | 385 | 0.96 | 405 | 1.01 | asm with multimap gDNA |
| cdsasm | 295 | 273 | 0.92 | 287 | 0.97 | asm with CDS-mapped gDNA |
| CDSann | 51 | 50 | 0.98 | 53 | 1.03 | asm with CDS annotations |
| TEann | 1476 | 1454 | 0.99 | 1531 | 1.04 | asm with Transposons |
| RPTann | 280 | 274 | 0.98 | 289 | 1.03 | asm with simple Repeats |
| NOann | 1284 | 1308 | 1.02 | 1377 | 1.07 | asm without annotations |

Total span=3117.2 Mb, covspan=3061.3, gaps=0 Mb for assembly human22ttot\_chr Genome Size Est= 3444.2-3627.0 Mb (Nread), 3368.5-3547.3 Mb (Maprd) for Size=LN/Cm, Cm=56.50,59.5, N\_reads=819708515, N\_maprd=97.8%, L=250

## Discussion & Conclusions

*The Goldilocks problem in genome reconstruction*. Despite results given on whole genome sizes, the focus of this new measurement tool is at detail levels of DNA coverage of genes and major components in chromosome assemblies. Those levels provide genomicists with a better link to biological knowledge for interpreting accuracy and completeness of gene and chromosome set reconstructions. As readers can guess, and indicated in Figure 9b with Daphnia assemblies, a “just-right” whole genome measure can be very wrong in details, but averages too-cold and too-hot to a false correctness.

The best example for Gnodes use may be comparing DNA depth deviations and major components of Arabidopsis chromosome assemblies (Figure R6.1). This offers graphic evidence of over- and under-assembly locations, and coincidence with centro- and telomeric repeat spans. Likewise chromosome depth plots for other species assemblies compared here offer insights and directions on where assemblies can be improved.

The recent shift to long read DNA for assembly can produce more complete chromosomes, yet they are often still incomplete by DNA content analyses and molecular size measures. Long read assemblies may even be notably less complete than prior Illumina or Sanger read assemblies (e.g. Daphnia and cucumber reported here, birds reported in Peona et al 2018). Assembly methods, as well as effort, and quality of DNA samples, have roles in this. As found with Gnodes and flow cytometry, recent long read assemblies for Arabidopsis, Daphnia species, eel, cucumber and sugar beet are incomplete in measurable ways. But the same DNA samples can be re-assembled with other methods to a fuller representation of that DNA.

A case in point is the Arabidopsis data set of Naish et al. (2021): their published assembly adds centromere spans to euchromatic arms using a mix of long read sets and assembly methods (Flye assembler) for 134 Mb total, but still lacks 10% of DNA content (Table R6.1, at21ncbi). A re-assembly of their pacbio hifi DNA with Canu assembler brings this total to 146 Mb, adding at telomeric ends and elsewhere, reducing the discrepancy to under 5% (Table R6.1, at22canu). Re-assembly of their nanopore data with Canu at 141 Mb is less complete. The majority of content in these assemblies are nearly same, comprising 120 Mb of euchromatic regions. The pacbio-hifi data add many short, repetitive spans that agree with DNA depth measured from Illumina data, most in telomeric repeats of Chrs 2 and 4 (Figure R6.1).

Gene coverage tables provide evidence for reconstructing more accurate gene sets, indicating status as unique, duplicated, or aberrant for a given DNA sample, with copy number measures. When combined with measures of genes located on chromosome assemblies, these also indicate where assemblies may be missing genes or parts, unique and duplicated, or have too many copies. The DNA depth deficit plot of Gnodes brings together chromosome and gene measures, to adequately, if imperfectly, summarize discrepancies in the observed assembly with DNA sample contents.

Whole genome measures, of size and other factors, are statistically complex, prone to errors of sometimes false assumptions, and should be supported by biological evaluation of details as much as possible. For instance, the K-mer estimators rely on an assumption of Poisson distribution for the depth of small, equalsize fragments of DNA [Li & Waterman 2003]. There are methodological and biological factors that invalidate these assumptions, but detecting and avoiding those violations is difficult. DNA depth based on unique genes coverage is based on a biological premise that is testable, that is are these genes unique in phylogenetic samples? If so, they represent spots of fixed DNA depth under the methodological premise of equal random samples from that genome. Results from Gnodes measures on unique genes produce tight, normally distributed values of depth.

Gnodes produces several genome size estimates (GSE), one of which is closer to true:

A. GSE for all reads is closest to true size of DNA sample when assumptions hold, i.e. no significant contamination or read error, same number of copies in all chromosomes. All reads are also the basis for K-mer methods and Gnodes estimate from genes alone.
B. GSE for mapped reads, which removes putative contamination and sample errors, if those exist in DNA but not assembly, but includes effect of incomplete assemblies.
C. Low and high measures of unique gene Cm, usually from gene-CDS mapped DNA (lower Cm, higher GSE), and from chromosome-assembly mapped DNA at unique gene loci (higher Cm, lower GSE). With clean data, these are nearly same values. Factors affecting these include gene map effects from number and sizes of exons (not known in transcripts), and chromosome assembly accuracy at UCG loci. Where these Cm measures diverge, the mid-point may be a best single estimate.

Investigators using this tool will choose among these size estimates based on knowledge of their DNA samples, chromosome assemblies and gene set.

In these results, violations exist of assumptions of equal copy numbers (diploid or haploid) of chromosomes with constant DNA depth. Sampled DNA has multiple copies of non-nuclear genomes (chloroplasts especially), and variable copies of sex chromosomes. Chloroplast genomes in plants can exist in 10,000 copies, depending on tissue sample. Chloroplast DNA measured in Arabidopsis samples ranged from 15 Mb to 65 Mb, relative to 157 Mb of 1-copy nuclear DNA, and was excluded from calculations of nuclear genome size. mtDNA, though a fraction of that (∼1 Mb), was also excluded. A human female DNA sample had 10% cover depth for ChrY, similarly for chicken sex ChrW/Z, and lower ChrY depth in fruitfly.

*Which is too hot, which is too cold?* Current genome informatics in this area has two general approaches, one that accepts that assemblies and K-mer estimators are in truth smaller than flow cytometry measures, another that rejects these as incomplete, under-sized results. The K-mer methods appear to under-estimate duplicated DNA. Results given here recommend use of two or more approaches for validating chromosome assembly sizes and contents, and examination of discrepancies to resolve those.

EvidentialGene use for gene set reconstruction is increasing in biomedical genomics including human disease studies. A recent human disease study of a pathogenic amoeba (Phan et al. 2020) with Evigene led to discovery of therapeutic compounds targeting amoeba genes. A USDA resource for pig genes, relevant to agricultural and biomedical uses, benefits from Evigene (Dawson et al. 2020).

A remaining objective is full integration of this DNA (chromosomes) tool with RNA (genes) reconstruction provided by EvidentialGene. This is fairly straight-forward, as Gnodes outputs contain needed information. The value of accurate duplicate gene sets is clear from biological studies on the role of duplications in rapid evolution and adaptation to environmental, disease and other organismal needs. Current informatics for duplicated genes are often inadequate. Evigene is well suited to accurate recovery of these, and is used in studies of often-duplicated venom genes (Modahl et al. 2020; Hanf et al. 2020), plant disease resistance genes (Rao et al. 2019), animal receptor and other duplicated genes (Hearn et al. 2020), and other difficult areas of gene reconstruction.

## Acknowledgements

I thank the following for helpful comments and suggestions: Boas Pucker, James Pflug, Spencer Johnston, Ilia Leitch, and scientists involved in Daphnia genomics. I thank the following for providing shared computational resources in support of this work: Extreme Science and Engineering Discovery Environment (XSEDE, www.xsede.org), including Jetstream (jetstream-cloud.org) and San Diego Supercomputer Center, Indiana University Pervasive Technology Institute (pti.iu.edu), and National Center for Genome Analysis Support (ncgas.org).

## Note

This is a draft document for public review, comments, and to facilitate use of Gnodes. Please comment to me (gilbertd at indiana.edu), or EvidentialGene’s public discussion (, https://groups.google.com/g/evidentialgene). Find EvidentialGene project at http://eugenes.org/ and http://sourceforge.net/projects/evidentialgene/

## Algorithm and Methods

The basic algorithm of Gnodes is (a) align DNA reads to gene and chromosome assembly sequences, recording all multiple and unique map locations, (b) tabulate DNA cover depth at each sequence bin location (100 bp bins) for multiple and unique mappings, (c) measure statistical moments of coverage depth per item (itemized by several categories of gene-cds, chr assembly, transposons and repeat annotations, duplicate and unique mappings, (d) summarize coverage and annotation tables, at chromosome-assembly and gene levels. The statistical population of coverage is non-normal, so that median, average, skew and other statistics are calculated to approach precise and reliable measures. Supplemental data include documentation on Gnodes methods, and software use results.

Effort and extensive testing has been used to implement details of these measures to achieve reliability with a variety of species genome data sets. For instance, the gene coverage depth algorithm (gnodes_sam2genecov.pl) examines per-base coverage, recording values over the gene-cds sequence span, and assesses coverage for span-completeness, span-unevenness, unique and duplicate mappings in the coding span, and median, average, skew values to classify each gene’s DNA coverage. The chromosome coverage depth algorithm (gnodes_sam2covtab.pl) includes measure of DNA reads that map to both chromosome and gene sequences (optionally adding transposon sequences), and incorporates annotations that partition the assembly into content types.

### GNODES PIPELINE STEPS

STEP 1. Annotations of Genes, TE on chr_assembly sh_repeatmask() if($doREPMASK and not $teseq) sh_buscoscan(); sh_annot($chrasm,$cdsseq,$teann,$crclassf,$buscout);

uses EVIGENES/genoasm/gnodes_annotate.pl, includes chr x cds align via BLASTN

STEP 2. DNA mapping

($gnbam)= sh_dnamap($cdsseq,$reads,@morereads); ($tebam)= sh_dnamap($teseq,$reads,@morereads); ($crbam)= sh_dnamap($chrasm,$reads,@morereads);

STEP 3. Tabulate read coverages

($genescovtab)= sh_sam2genecov($gnbam, ..) ;

uses EVIGENES/genoasm/gnodes_sam2genecov.pl

($cdstab) = sh_sam2covtab(‘cds’, $gnbam, ..);

($tetab) = sh_sam2covtab(‘te’, $tebam, ..); ($chrcovtab) = sh_sam2covtab(‘chr’, $crbam, ..);

uses EVIGENES/genoasm/gnodes_sam2covtab.pl

Note: chr.bam and cds.bam must have same reads mapped in same order, gnodes_pipe does this. STEP 4: Summarize

($sumcds)= sh_covsum($cdstab, $cdsid, $cdsid, ““, $crclassf, $genescovtab, ..);

($sumout)= sh_covsum($chrcovtab, $asmid, $intitle, $anntable, $crclassf, $genescovtab);

uses EVIGENES/genoasm/gnodes_covsum.pl

($gsumout)= sh_sumgenescov($genescovtab, $chrgenetab, $asmid, ..); uses EVIGENES/genoasm/gnodes_sumgenecov.pl

### GNODES USAGE

~~~
$evigene/scripts/genoasm/gnodes_pipe.pl \
 -title output_prefix \
 -chr chr_assembly.fasta -cds cds.fasta \ # required
 -reads SRRnnnn_1.fastq # required, may be many files
 eg.-reads SRR01_[12].fastq SRR02_[12].fastq
~~~

options:

~~~
 -cds may be replaced with -mrna gene_transcripts.fasta
 -asmid chr_assembly : usually chr_assembly name as in -chr fasta
 -metadata genomes.metadata : information for each asmid (below)
 -idclasses cdste.idclass : table of classes per gene cds or transposon (te) identifier
 -te te.fasta : optional input transposon.fa sequences, or use RepeatMasker
 -ncpu 24 : number-of-threads to use
 -maxmem 120gb : maximum computer memory, 120 Gb is likely sufficient for large genomes like human.
 -RDMAPPER bwa-mem2|minimap2 : one of a few read-mapper applications that are known to gnodes_pipe.
~~~

**gnodes_pipe.pl** generates a shell script **run_gnodes.sh**, using input file names and options. The run_gnodes.sh calls several component Evigene scripts with those inputs, suited for cluster computers. gnodes_pipe.pl does no genome computations, and is safe to run on non-cluster machine.

**gnodes_setup.sh** : Settings for compute cluster and Unix system to be included at top of run_gnodes.sh

### GNODES REQUIRED SOFTWARE

EvidentialGene, evigene21jul30.tar or later, from http://eugenes.org/EvidentialGene/other/evigene_old/ Read mapper, bwa-mem2 or minimap2 currently, from https://github.com/bwa-mem2/bwa-mem2/,

https://github.com/lh3/minimap2/

samtools from https://sourceforge.net/projects/samtools/

NCBI BLAST, from ftp://ftp.ncbi.nlm.nih.gov/blast/

BUSCO, run_BUSCO.py and hmmer software, Orthodb data for species, version 3 okay, other versions need test, from https://busco.ezlab.org/, https://gitlab.com/ezlab/busco/tree/3.0.2/

RepeatMasker to find transposons/repeats, or transposon sequences, from http://www.repeatmasker.org/ R statistical package is used to draw summary, chromosome plots.

### METADATA FOR ASSEMBLIES

A text file with these key=value entries is needed for full use of Gnodes, with each entry for an assembly id (asmid=) followed by entries of the other keys. There may be many asmid records in the metadata file.

asmid= assembly ID, often name of chrseq.fasta file

flowcyto= flow cytometry genome size

asmtotal= Chr assembly size, for _total summary

asmname= Label for asmid

species= Genus_species

buscodb= OrthoDB database name for BUSCO calculation

rmaskdb= RepeatMasker database

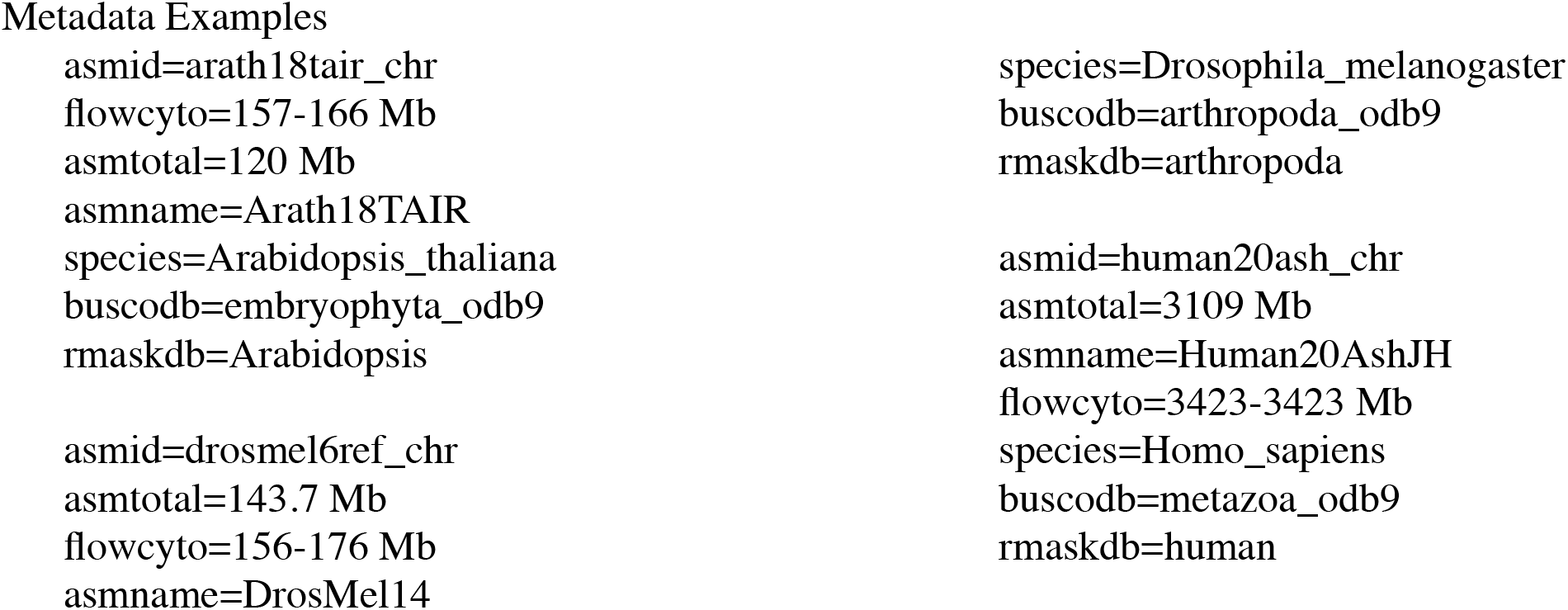

### EXAMPLE PLANT

DNA samples:

a. SRR10178325 of Bioproject PRJNA574113, Illumina HiSeq, 6G bases, of Arabidopsis thaliana leaf whole genome sequence of homozygous of Col-0 strain
b. SRR3703081 of PRJNA314706, 4.8G bases, F1 heterozygote of Col-0 and Cvi-0 strains.

CDS gene sequences used

a. arath18tair1cds.fa, TAIR 2018, chr-asm modelled, isoform 1 only
b. arath17evg5ico1cds.fa, Evigene RNA-assembled 2017, isoform 1 only

CDS set b is used for assembly comparisons as it is independent of the chr-assemblies, has more unique conserved genes, though is otherwise similar to chr-modeled gene set. Gnodes gene results are similar though the alternate 2020 Max assembly has slightly greater recovery of Evigene set than that modeled on TAIR 18 chr assembly.

Chromosome assemblies tested:

a. arath18tair_chr, Reference chr-assembly, Total length: 119.7 Mb, from ftp://ftp.ncbi.nlm.nih.gov/genomes/all/GCF/000/001/735/GCF_000001735.4_TAIR10.1/
b. arath20max_chr, Alternate chr-assembly, Total length: 130.2 Mb, Submitter: Max-Planck Institute For Developmental Biology, 2020/09/12, from ftp://ftp.ncbi.nlm.nih.gov/genomes/all/GCA/904/420/315/GCA_904420315.1_AT9943.Cdm-0.scaffold

Gnodes invocations:

~~~
pt=arath18tair_chr; pt=arath20max_chr;
$evigene/scripts/genoasm/gnodes_pipe.pl -title ${pt}8evg -chr $pt.fa -cds arath17evg5ico1cds.fa -sumdata arath20asm.metad -ncpu 32 -maxmem 180gb -RDMAPPER bwa-mem2 -reads readsf/SRR10178325_[12].fastq.gz
~~~

### EXAMPLE FLY

DNA sample: SRR11460802 of PRJNA618654 VetMedUni Vienna, Drosophila melanogaster, 2020-11-25, Illumina HiSeq, 5.7G bases.

CDS gene sequences: dromel6rel_cds.fa, dromel ncbi refseq of rel. 6, isoform=1 only Chromosome assemblies tested:

a. drosmel6ref_chr, Reference chr-assembly, Total sequence length: 143.7 Mb, BioProject: PRJNA13812; GenBank accession: GCA_000001215.4; Name: Release 6 plus ISO1 MT; Submitter: The FlyBase Consortium/Berkeley Drosophila Genome Project/Celera Genomics, 2014-08-01

b. drosmel20pi_chr, Alternate chr-assembly, Total sequence length: 167.8 Mb; BioProject: PRJNA618654; GenBank accession: GCA_015852585.1; Submitter: VetMedUni Vienna, 2020/12/08 Gnodes invocations:

~~~
pt=drosmel6ref_chr; pt=drosmel20pi_chr;
$evigene/scripts/genoasm/gnodes_pipe.pl -title $pt -chr $pt.fa -cds dromel6rel_cds.fa-metadata drosmelchr.metad -ncpu 24
-maxmem 128gb -reads readsf/
SRR11460802_[12].fastq
~~~

### EXAMPLE HUMAN

DNA sample: SRR12898282 of PRJNA200694, HiSeq2500_LAB01_Mother_REP01, Illumina HiSeq, 132.8G bases, 2020-10-26.

CDS gene sequences: human18nc_cds1t.fa, human ncbi refseq of 2018, isoform=1 only. Chromosome assemblies tested:

a. human19grc_chr, Reference chr-assembly, Total sequence length: 3,099 Mb, Name: GRCh38.p13, BioProject PRJNA31257, GenBank accession: GCF_000001405.39 ; Submitter: Genome Reference Consortium, 2019/02/28

b. human20ash_chr, Alternate chr-assembly, Total sequence length: 3,109 Mb, Name: Ash1.7, BioProject PRJNA607914, GenBank accession: GCA_011064465.1, Submitter: Johns Hopkins University, 2020/03/04

Gnodes invocations:

~~~
pt=human19grc_chr; pt=human20ash_chr;
$evigene/scripts/genoasm/gnodes_pipe.pl -title $pt -chr $pt.fa -cds
human18nc_cds1t.fa -metadata human20asm.metad -ncpu 32 -maxmem 240gb -RDMAPPER bwa-
mem2 -reads readsf/SRR12898282_[12].fastq.gz
~~~

## Supplemental data

### gnodes_docsup : detail tables supporting paper text, tables, and figures

S1_arath17evg5ico1cds_SRR10178325_b2.genexcopy.txt

Table of Gnodes genes DNA depth and copy number output, as abstracted in Table R1.1. Genes DNA coverage table example of Arabidopsis thaliana (arath17evg gene set, SRR10178325 DNA reads). The Uniq class column identifies 4 types: uniq, dupx (duplicated), dups (partial duplicate), skew (uneven coverage), and zero (coverage below reliability). Columns Glen, Nread are length of coding sequence, and number of reads mapped. tCopy is total copy estimate, C.M is coverage depth measure, Merr is map error percentage, C.nz is a tuple with average, median, length and percent covered span. The table top summary has unique gene cover depth measures, and genome size estimate from those and the read length and number in DNA sample.

S2_gnodes_allsppucg_combotab8i.txt

Table for Figures R1.1, R1.2, containing data of 13 species, DNA samples, genome sizes and data from Gnodes analyses. Columns are Genome_Name (species, assembly), SRA_Reads, G_fcyto (GSE of flow cytometry), G_asm (GSE of assembly), G_ucg (GSE of Gnodes UCG), C.M.mdn_sem (Gnodes Cucg median, sem), Nr_tot,Lrd,ucg%,cds%,err%, a tuple with N-reads, length of reads, percent of reads in UCG, CDS, read errors, and Info on species data.

S3_gnodes_asfcuckh_tab8i.txt

Table for Figures R1.1, R1.2, containing data of 13 species, with GSE of K-mer methods, Gnodes UCG, flow cytometry, assembly size.

S4_arath18tair_nopcr36ecotypes_gnodes_table.txt

full table for items referred to in Table R8.1, Figure R8.1 and text, comprises data from 36 DNA samples, PCR-Free Illumina reads, of Arabidopsis thaliana ecotypes published by Van De Weyer et al 2019. Table columns are NoPCR_ID for read set in SRA; Etype, Lon, Lat, Country are ecotype ID, longitude, latitude, country tag; FC16s, FC16et are closest flow-cytometry size measure and ecotype ID from Cao et al 2011; GScope, findGSE are k-mer method genome size estimates of DNA; LNCmap, LNCtot are Gnodes size estimates (mapped reads, total reads - contaminants), CU is Gnodes DNA depth at unique genes, allm, uniqm, dupm, CDSa, TEa, RPTa, NOa are Gnodes partition sizes in Mb, xUnixm, xNoa, xCDSa are xCopy values of those partitions; CONTAM is removed contaminant (plastid) size, covspan is assembly coverage span; Maprd, pMap are mapped read count, percentage; tabdate date of tabulation.

NoPCR_ID Etype, Lon Lat Country FC16s FC16et GScope findGSE SRA_Reads LNCmap LNCtot CU allm uniqm dupm CDSa TEa RPTa NOa xUniqm xNOa xCDSa CONTAM covspan Maprd, pMap tabdate ERR2178830 9869,Moj-0 -5.28 36.76 ESP NA 0 124.5 128.2 ERR2178830ab 161 168 82.2147 67 33 40 13 16 42 1.06 1.10 1.01 32 115.2 63278774,96.2 2022_Mar_21

S5_gnodes22species_metad.txt

Table of sources for chromosome assemblies and genes.

S6_gnodes_about_pb.txt

Documentation describing Gnodes code, usage, required software, pipeline steps, outputs, metadata required and example plant and animal runs.

S7_read_mappers_compared.txt

Table comparing read map software, bwa-mem2 and minimap2, effects on Gnodes statistics. bwa maps ∼0.5% more reads than minimap (pMap), for default options, with elevated error rate (Merr).

### gnodes_doctabs : gnodes outputs per species, assembly

~~~
9.4M Nov 12 arath20asm_test8tf.tar.gz
  for arath18tair and arath20max assemblies,
  refseq and evigene gene sets, with SRR10178325 DNA sample
18M May 4 at22vlr_ncbi_gnodes.tar.gz
 for at22vlr_ncbi_chrs assembly,
 refseq and evigene gene sets, with SRR10178325 DNA sample
3.4M May 4 at22vcanu2d_gnodes.tar.gz
 for at22vcanu2pbhi_chrs assembly,
 refseq and evigene gene sets, with SRR10178325 DNA sample
5M May 3 arath18tair_nopcr36ecotypes_gnodes.tar.gz
 for arath18tair assembly, refseq genes, with PCR-Free DNA samples of 36 ecotypes
11M May 1 beet22gnodes.tar.gz
 beet18usda assembly, beet15 refseq genes, SRR6305245 DNA
73M Nov 12 chick19nc_test8jf.tar.gz
 chick19nc_chr assembly, refseq genes, SRR3954707 DNA
11M May 1 corn20mgd_gnodes8lf.tar.gz
 zmays20MGD5 assembly, refseq and evigene genes, ERR3288215,6? DNA
21M May 4 cucum19cgi_test8jf.tar.gz
 cucum19cgi assembly, refseq genes, SRR11300859 DNA
33M Nov 14 dromel20asm_test8tf.tar.gz
 drosmel6ref and drosmel20pi assemblies, refseq genes, SRR11460802 DNA
7.8M May 1 eel22gnodes.tar.gz
 eueel21inra_chrs assembly, refseq genes, SRR5235521,2,3? DNA
146M May 4 pig20asm_test8jf.tar.gz
 pig11 chr assembly, evigene genes, SRR4341337 DNA
287M Nov 12 human19grc_test8jq.tar.gz
 human GRC assembly, SRR12898282 HG004 DNA sample
365M Apr 26 human22tt_gnod741f.tar.gz
 human T2T assembly, SRR3189741 CHM13 DNA sample
7.2M May 4 human22tt_gnod282f.tar.gz
 human T2T assembly, SRR12898282 HG004 DNA sample
~~~

### gnodes_newasm: species chromosome assemblies produced for this paper

*Arabidopsis thaliana*

at22vcanu2pbhi_chrs, for Table R6.1 AT3a3, Figure R6.1, text, at22vcanu2d_gnodes from pacbio hifi DNA SRR16841688, canu2.2 assembler, run_gasm_canu_arath22vlr2b.sh at22vcanu3nano_chrs, for text,

from nanopore DNA SRR16832054, canu2.2 assembler, run_gasm_canu_arath22vlr3n.sh Assembled contigs were mapped to at22vlr_ncbi_chrs assembly to produce pseudo-chromosomes.

*Daphnia magna* : to be added

*Daphnia pulex* : to be added

## References

Adams, M, J McBroome, N Maurer, E Pepper-Tunick, N F. Saremi, R E. Green, C Vollmers and R B. Corbett-Detig (2020). One fly–one genome: chromosome-scale genome assembly of a single outbred Drosophila melanogaster. Nucleic Acids Research, 48(13) e75; doi: 10.1093/nar/gkaa450

Arabidopsis Genome Initiative (2000). Analysis of the genome sequence of the flowering plant Arabidopsis thaliana. Nature 408: 796–815.

Bennett, MD, IJ Leitch, HJ Price and JS Johnston (2003) Comparisons with Caenorhabditis (100Mb) and Drosophila (175Mb) using flow cytometry show genome size in Arabidopsis to be 157Mb and thus 25% larger than the Arabidopsis genome initiative estimate of 125Mb. Ann. Botany, 91, 547–557 doi:10.1093/aob/mcg057

Cao J, K Schneeberger, S Ossowski, T Gunther, S Bender, J Fitz, D Koenig, C Lanz, O Stegle, C Lippert, X Wang, F Ott, J Muller, C Alonso-Blanco, K Borgwardt, K J Schmid & D Weigel (2011). Whole-genome sequencing of multiple Arabidopsis thaliana populations. Nature Genetics, 28 August 2011; doi:10.1038/ng.911

Chin, C-S, P Peluso, F J. Sedlazeck, M Nattestad, G T. Concepcion, A Clum, C Dunn, R O’Malley, R Figueroa-Balderas, A Morales-Cruz, G R. Cramer, Massimo Delledonne, C Luo, J R. Ecker, D Cantu, D R. Rank, and M C. Schatz (2016). Phased Diploid Genome Assembly with Single Molecule Real-Time Sequencing. Nat Methods. 13(12): 1050–1054. doi:10.1038/nmeth.4035

Cook, R.K., Christensen, S.J., Deal, J.A. et al. The generation of chromosomal deletions to provide extensive coverage and subdivision of the Drosophila melanogaster genome. Genome Biol 13, R21 (2012). doi: 10.1186/gb-2012-13-3-r21

Desvillechabrol, D., Bouchier, C, Kennedy, S and Cokelaer, T (2018). Sequana coverage: detection and characterization of genomic variations using running median and mixture models. Gigascience, 7(12), p.giy110. doi: 10.1093/gigascience/giy110

Dolez J & J Greilhuber (2010). Nuclear Genome Size: Are We Getting Closer? Cytometry; doi: 10.1002/cyto.a.20915

Gilbert, DG (2013). Gene-omes built from mRNA seq not genome DNA. 7th annual Arthropod Genomics Symposium. Notre Dame. doi:10.7490/f1000research.1112594.1

Gilbert, DG (2019). Genes of the Pig, Sus scrofa, reconstructed with EvidentialGene. PeerJ 7:e6374; doi:10.7717/peerj.6374

Gregory, TR. (2017). Animal Genome Size Database. http://www.genomesize.com.

Hanf, Z. R., & Chavez, A. S. (2020). A comprehensive multi-omic approach reveals a relatively simple venom in a diet generalist, the northern short-tailed shrew, Blarina brevicauda. Genome Bio. and Evo. doi: 10.1093/gbe/evaa115

Hearn J, J Clark, PJ. Wilson, and TJ. Little (2020). Daphnia magna modifies its gene expression extensively in response to caloric restriction revealing a novel effect on haemoglobin isoform preference. bioRxiv; doi: 10.1101/2020.05.24.113381

Hozza, M, T Vinar, and B Brejova (2015). How big is that genome? Estimating genome size and coverage from k-mer abundance spectra, pp. 199–209 in String Processing and Information Retrieval, edited by C. Iliopoulos, S. Puglisi, and E. Yilmaz. Lecture Notes in Comp Sci., Springer Intl. Pub. [CovEST]

International Human Genome Sequencing Consortium (2004). Finishing the euchromatic sequence of the human genome. Nature 431:931–945.

Jansen, H.J., Liem, M., Jong-Raadsen, S.A., Dufour, S., Weltzien, F.-A., Swinkels, W., Koelewijn, A., Palstra, A.P., Pelster, B., Spaink, H.P., et al. (2017). Rapid de novo assembly of the European eel genome from nanopore sequencing reads. Sci. Rep. 7, 1–13. doi: 10.1038/s41598-017-07650-6

Ji L., T Sasaki, X Sun, P Ma, Z A. Lewis and R J. Schmitz (2014). Methylated DNA is over-represented in whole-genome bisulfite sequencing data. Front. Genet., 21 October 2014; doi: 10.3389/fgene.2014.00341

Kim J, C Lee, B J Ko, D Yoo, S Won, A Phillippy, et al. (2021). False gene and chromosome losses affected by assembly and sequence errors. bioRxiv 2021.04.09.438906; doi: https://doi.org/10.1101/2021.04.09.438906

Lander, E S, and M S Waterman (1988). Genomic mapping by finger-printing random clones: A mathematical analysis. Genomics 2: 231–239. doi:10.1016/0888-7543(88)90007-9 [G=L*N/C]

Long Q, F A Rabanal, D Meng, C D Huber, A Farlow, A Platzer, Q Zhang, B J Vilhjalmsson, A Korte, V Nizhynska, V Voronin, P Korte, L Sedman, T Mandakova, M A Lysak, U Seren, I Hellmann, and M Nordborg (2013). Massive genomic variation and strong selection in Arabidopsis thaliana lines from Sweden. Nat Genet. 45(8): 884–890. doi:10.1038/ng.2678.

McGurk M P, A-M Dion-Cote, D A Barbash (2021). Rapid evolution at the Drosophila telomere: transposable element dynamics at an intrinsically unstable locus, Genetics, 217(2), iyaa027; doi: 10.1093/genetics/iyaa027

Modahl C.M., Durban J., Mackessy S.P. (2020). Exploring toxin evolution: venom protein transcript sequencing and transcriptome-guided high-throughput proteomics. In: Priel A. (ed) Snake and Spider Toxins. Methods in Molecular Biology, vol 2068. Humana, New York, NY; doi: 10.1007/978-1-4939-9845-6_6

Naish M, M Alonge, P Wlodzimierz, A J. Tock, B W. Abramson, A Schmucker, T Mandakova, B Jamge, C Lambing, P Kuo, N Yelina, N Hartwick, K Colt, L M. Smith, J Ton, T Kakutani, R A. Martienssen, K Schneeberger, M A. Lysak, F Berger, A Bousios, T P. Michael, M C. Schatz, I R. Henderson (2021). The genetic and epigenetic landscape of the Arabidopsis centromeres. Science 374, eabi7489. doi: 10.1126/science.abi7489

Nurk S, Koren S, Rhie A, Rautiainen M, et al (2022). The complete sequence of a human genome. Science 376, 44– 53; doi: 10.1126/science.abj6987

Pellicer J, Ilia J Leitch (2019). The Plant DNA C-values database (rel 7.1): an updated online repository of plant genome size data for comparative studies. New Phytologist, 226(2): 301–305; doi: 10.1111/nph.16261

Peona V, Weissensteiner MH, Suh A (2018). How complete are “complete” genome assemblies? An avian perspective. Mol Ecol Resour. 18(6):1188–1195. doi: 10.1111/1755-0998.12933.

Pflug JM, V R Holmes, C Burrus, JS Johnston, and DR Maddison (2020). Measuring Genome Sizes Using Read-Depth, k-mers, and Flow Cytometry: Methodological Comparisons in Beetles (Coleoptera). Gen.Gen.Gen., 10:3047-3060; doi: 10.1534/g3.120.401028

Phan, I Q, C A Rice, J Craig, R E Noorai, J McDonald, S Subramanian, L Tillery, L K Barrett, V Shankar, J C Morris, W C Van Voorhis, D E Kyle, P J Myler (2020). The transcriptome of Balamuthia mandrillaris [pathogenic amoeba] trophozoites for structure-based drug design. bioRxiv doi:10.1101/2020.06.29.178905; NIH dataset, doi: 10.35092/yhjc.12478733.v1

Pucker, B (2019). Mapping-based genome size estimation [MGSE]. bioRxiv, doi: 10.1101/607390

Rao T B, R Chopperla, R Methre, et al (2019). Pectin induced transcriptome of a Rhizoctonia solani strain causing sheath blight disease in rice reveals insights on key genes and RNAi machinery for development of pathogen derived resistance. Plant Molecular Biology; doi: 10.1007/s11103-019-00843-9

Rhie A, McCarthy SA, Fedrigo O, Damas J, Formenti G, Koren S, et al. (2021). Towards complete and error-free genome assemblies of all vertebrate species. Nature, 592(7856):737–746. doi: 10.1038/s41586-021-03451-0.

Simao FA, Waterhouse RM, Ioannidis P, Kriventseva EV, Zdobnov EM (2015). BUSCO: assessing genome assembly and annotation completeness with single-copy orthologs. Bioinformatics;31(19):3210–2. doi: 10.1093/bioinformatics/btv351. PMID: 26059717.

Sun, H, J Ding, M Piednoël, and K Schneeberger (2017). findGSE: estimating genome size variation within human and Arabidopsis using k-mer frequencies. Bioinformatics 34: 550– 557.

Van de Weyer AL, Monteiro F, Furzer OJ, et al. (2019). A Species-Wide Inventory of NLR Genes and Alleles in Arabidopsis thaliana. Cell. 178(5):1260-1272.e14. doi: 10.1016/j.cell.2019.07.038.

Vurture, GW, Sedlazeck, FJ, Nattestad, M, Underwood, CJ, Fang, H, Gurtowski, J, Schatz, MC (2017). GenomeScope: fast reference-free genome profiling from short reads. Bioinformatics 33: 2202–2204. doi: 10.1093/bioinformatics/btx153

Yoon S, Z Xuan, V Makarov, K Ye and J Sebat (2009). Sensitive and accurate detection of copy number variants using read depth of coverage. Genome Res. 19:1586–1592; doi: 10.1101/gr.092981.109

